# Rapid odorant metabolism organizes identity- and timing-based odor representations by olfactory bulb inputs and outputs

**DOI:** 10.64898/2026.07.01.735924

**Authors:** Elvis F. Acquah, Madison A. Herrboldt, Joel Hallkaj, Mona A. Marie, Zhenxing Wu, Jeanne Chaloyard, Isabelle Andriot, Jean-Marie Heydel, Kai Zhao, Hiroaki Matsunami, Matt Wachowiak

## Abstract

Sensory neurons encode information about external stimuli in the form of stimulus-specific patterns of activity across their population. In the mammalian olfactory system, olfactory sensory neurons (OSNs) encode odor identity with odorant-specific combinatorial patterns that are temporally structured by inhalation. We investigated the stimulus features that determine these inhalation-linked patterns, leveraging receptor- and functionally-defined OSN populations in awake mice. We found that both the chemical tuning and temporal dynamics of many odorant-evoked responses deviate from prevailing models of receptor-ligand binding and sensitivity-based timing relationships. These deviations were well-explained by rapid metabolism of odorants within the olfactory mucosa, which generates secondary odorants that activate additional OSNs within a single breath. This process fundamentally reshapes odor representations at naturally-occurring concentrations and timescales relevant to perception. Importantly, we found that timing features robustly discriminate inhaled from metabolism-generated odorants, and that these timing differences persist at the level of olfactory bulb output. Our results suggest a novel role for inhalation-linked timing in odor coding - to disambiguate inhaled odorants from those generated internally - and raise the possibility that the nervous system may differentially process external and internally-sourced olfactory stimuli on the basis of their temporal dynamics.

## Introduction

In the mammalian olfactory system, inhalation delivers odorants to the olfactory epithelium and activates olfactory sensory neurons (OSNs) with diverse temporal patterns. Inhalation-linked timing features carry information about odorant identity and intensity (Spors et al., 2006; Bathellier et al., 2008; Cury and Uchida, 2010; Shusterman et al., 2011), and there is strong evidence that the sequence of activation of OSNs or mitral/tufted (MT) cell outputs from the olfactory bulb (OB) can impact odor perception (Smear et al., 2013; Rebello et al., 2014; Wilson et al., 2017; Chong et al., 2020). The determinants of diverse inhalation-linked temporal patterns and their role in mediating adaptive behavioral responses to olfactory stimuli remains an open question in understanding the neural basis of odor sensation.

A prominent model proposes that inhalation-linked timing reflects the relative sensitivity of an OSN population to an odorant, with the most-sensitive OSNs activated earlier and preferentially driving odor perception (Wilson et al., 2017). By this ‘primacy’ model (Giaffar et al., 2024), sensitivity-determined temporal patterning enables concentration-invariant representation of odor identity because the sequence of activity remains constant across concentration (Hopfield, 1995; Wilson et al., 2017). Sensitivity-timing relationships have not been systematically tested in vivo, partly because of the difficulty in defining the most-sensitive (i.e.,‘primary’) OSN population for a given odorant and the difficulty in probing functionally-defined OSN populations in vivo.

While sensitivity-based timing would seem to follow from first principles, the path from vapor-phase odorant to receptor-ligand binding in vivo is complex and involves multiple steps including a tortuous flow path through the nasal cavity, absorption into an aqueous mucus phase, partitioning into the underlying epithelial tissue and interactions with accessory proteins in the mucus where receptor-bearing cilia are located (Mozell, 1964; Yang et al., 2007; Lawson et al., 2012; Heydel et al., 2013). Any of these steps could perturb the kinetics and concentrations of odorants in a compound-specific manner during inhalation.

A particular interaction with the potential to alter sensitivity-timing relationships is the metabolism of odorants as they interact with nasal tissue. The nasal cavity is a vulnerable point of contact between the interior of an organism and potentially harmful chemicals in its environment. To combat this hazard, enzymes expressed in the nasal mucosa (epithelium + mucus) convert incoming chemicals to more water-soluble forms that are more easily removed from tissue (Heydel et al., 2013; Heydel et al., 2019). These ‘xenobiotic’ enzymes are among the highest-expressed proteins in the nasal epithelium (Ibarra-Soria et al., 2014) and many have been detected in the mucus surrounding OSN cilia (Olson et al., 1993).

Odorant metabolism has widely been considered a ‘housekeeping’ process with little impact on the encoding of odor identity. However, metabolites of inhaled odorants can themselves function as odorants and there is evidence in humans that odorant metabolism occurs within a single breath and shapes odor perception (Kornbausch et al., 2022; Robert-Hazotte et al., 2022; Shirai et al., 2023). In mice, a few odorant - enzyme interactions have been linked to activation of defined populations of OSNs and thereby to altered glomerular responses in the olfactory bulb and, potentially, altered odor perception (Nagashima and Touhara, 2010; Kida et al., 2018). Given the variety of enzymes present in the olfactory mucosa (Heydel et al., 2019) and the range of their chemical substrates and products, nasal metabolism has the potential to broadly impact odor representations.

Here, we sought to define the determinants of timing- and identity-based odor representations using functionally- and receptor-defined OSN populations for which high-potency odorants were known a priori and characterizing their chemical tuning properties and inhalation-linked response dynamics in awake mice. We found that, for major subsets of OSNs and major (i.e., commonly-encountered) chemical classes of odorants, the tuning of OSNs to chemical features as well as their inhalation-linked response dynamics were not explained by receptor-determined, sensitivity-based models. These deviations were well-accounted for by a model in which nasal enzymes rapidly produce odorous metabolites, activating metabolite-sensitive OSNs with a slight delay relative to inhaled odorants, but nonetheless within the same respiratory cycle. We present multiple lines of evidence supporting the conclusion that rapid nasal metabolism of odorants profoundly alters identity, chemotopic and timing-based representations of olfactory stimuli, potentially confounding the accurate encoding and recognition of external chemical signals. At the same time, we find that inhalation-linked timing features effectively disambiguate inhaled versus metabolism-generated odorants and that the delayed inputs characteristic of metabolite signals are preserved at the level of OB output. These results suggest an alternative function for perceptual primacy and timing in odor coding – namely, to disambiguate sensory signals arising from external versus internal odor sources.

## Results

### Odorant metabolism explains in vivo tuning of class I OR-expressing OSNs

We first considered evidence that odorant metabolism shapes the chemical tuning of OSNs in vivo, using a recent comprehensive characterization of OSN sensitivities to odorant chemical features at the level of their input to OB glomeruli (Burton et al., 2022). We observed that response maps of OSN input to dorsal OB glomeruli were nearly identical for low (pM - nM) concentrations of a given carboxylic acid and its aldehyde and ester derivatives (Fig. 1A; Supplemental Figure S1). Analysis of odorant co-tuning for carboxylic acid-sensitive glomeruli revealed that, despite co-tuning between acid, ester, and aldehyde derivatives, OSNs were nonetheless narrowly tuned to odorants *within* a functional group, responding selectively to a narrow range of carbon chain backbone structures (Fig. 1B).

**Figure 1.**
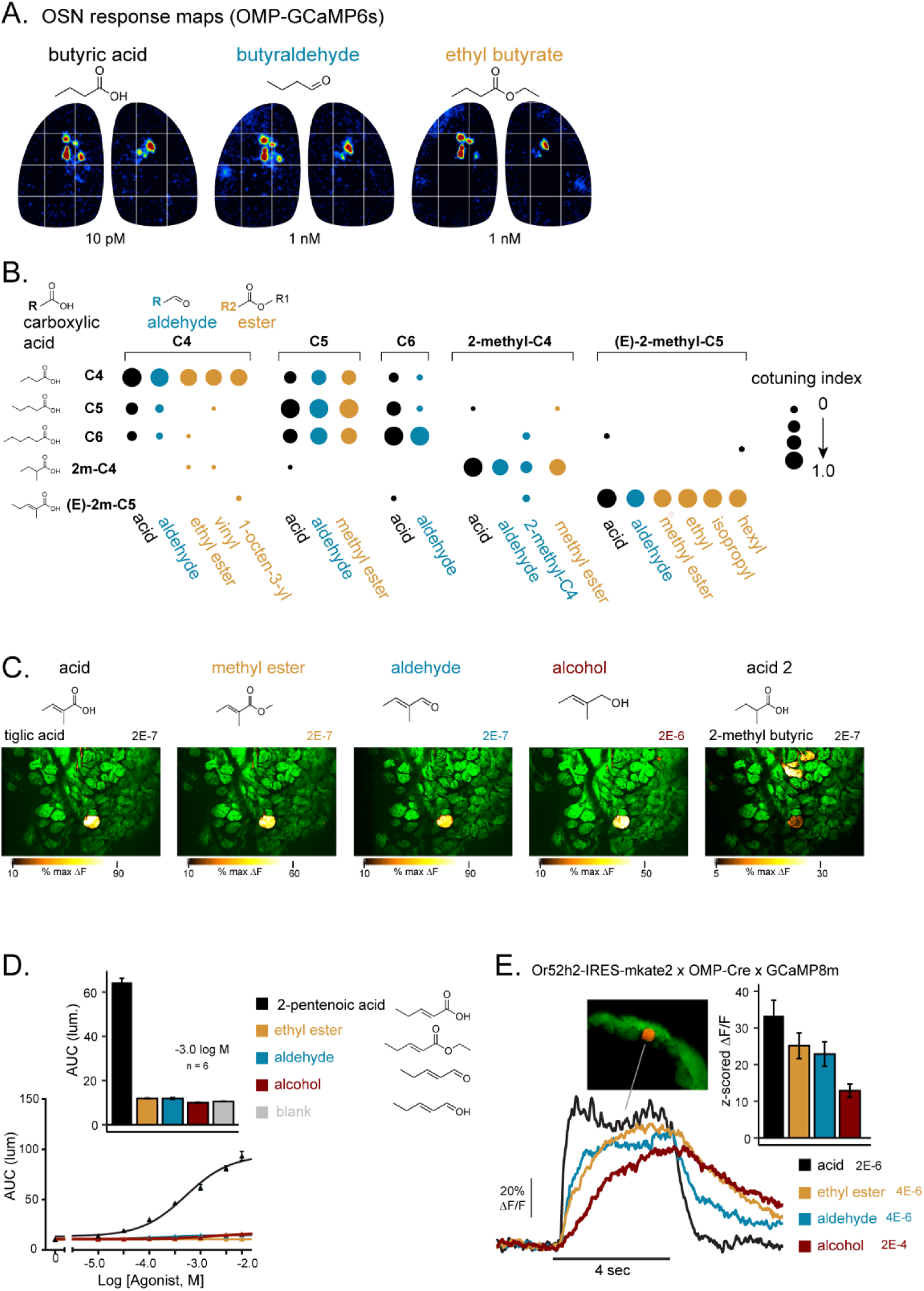
In vivo co-tuning of OSNs to carboxylic acids and their ester, aldehyde and alcohol derivatives is explained by odorant metabolism. **A.** Butyric acid and its aldehyde and ester derivatives elicit near-identical response maps of OSN input to dorsal OB glomeruli at estimated delivered concentrations ranging from 10 pM to 1 nM. Data from (Burton et al., 2022); OMP-IRES-tTA x tetO-GCaMP6s mouse. **B.** Dot plot of glomerular co-tuning across a larger panel of odorants, analyzed from (Burton et al., 2022), showing systematic co-tuning to acids, ester and aldehyde derivatives but selectivity across changes in carbon backbone structure. See Methods for co-tuning index definition. **C.** Potent and selective activation of OSN inputs to a functionally-identified glomerulus with primary sensitivity to tiglic acid by its ester, aldehyde and alcohol derivatives, imaged in the awake, head-fixed mouse (OMP-Cre x Tigre2-GCaMP8m). A structural variant, 2-methyl butyric acid (2-MBA) weakly activates this glomerulus strongly activates distinct glomeruli. Numbers indicate total odorant dilution (liquid dilution x vapor-phase dilution; see Methods). **D.** Selectivity of the class I odorant receptor Or52h2 to the carboxylic acid 2-pentenoic acid but not its ester, aldehyde or alcohol derivatives in a heterologous expression system. Left: Concentration-response plot for each odorant; Units expressed as area under the luminescence curve, AUC (see Methods). Right: Mean response (at 1E-3 M) for each odorant. ethyl ester: E-ethyl-pent-2-enoate; aldehyde: trans-2-pentenal; alcohol: trans-2-penten-1-ol. Error bars indicate SEM (n = 3 experiments per odorant, 3 technical replicates per experiment). **E.** Imaging OSNs expressing Or52h2 in vivo (tagged with mKate2 for visualization) reveals co-tuning to the ester, aldehyde and alcohol derivatives in the awake mouse. Inset, sample OB section showing Or52h2-tagged glomerulus (red) with native GCaMP fluorescence in OSNs (green). Left: Example trial-averaged traces (4 presentations) showing response to presentation of 2-pentenoic acid and the ethyl ester, aldehyde and alcohol derivatives. Right: Mean response amplitudes to each odorant. Error bars, SEM; n = 7 mice.

We confirmed this pattern of odorant tuning in awake mice expressing GCaMP8m in OSNs, focusing on four functionally-identified OSN populations defined by their characteristic inputs to glomeruli in stereotyped locations (Burton et al., 2022) (glomeruli 10, 23, 5, 11). Each of the four glomeruli were singularly activated by estimated picomolar concentrations of a particular carboxylic acid (Supp. Fig. S1) and also by its derivative ester and aldehyde; two of the glomeruli were additionally tested with and responded robustly to the *alcohol* derivative (Fig. 1C, Supp. Fig. S1).

These tuning properties are surprising given that a recent characterization of a human class I OR (OR51E2) identified a structural basis for high selectivity to carboxylic acids (Billesbølle et al., 2023). We confirmed this selectivity by expressing a different class I OR, mouse Or52h2, also known as MOR31-7 and Olfr649, in a heterologous expression system (Saito et al., 2009) and challenging it with odorants in vitro. Or52h2 responded sensitively to its preferred acid ligand, 2-pentenoic acid, but *not* its ester, aldehyde or alcohol derivatives (Fig. 1D). We next generated a mouse line tagging Or52h2-expressing OSNs (Or52h2-IRES-mKate2), crossed this line into an OMP-Cre x TIGRE2-GCaMP8m strain, and performed two-photon imaging from Or52h2 OSN terminals in its dorsal glomerulus in awake mice (n = 7). In this setting, Or52h2-expressing OSNs responded robustly to 2-pentenoic acid as well as the derivative ester, aldehyde, and alcohol (Fig. 1E).

Together, these results suggest that the tuning of acid-sensitive OSNs to esters, aldehydes, and alcohols is not a function of the OR itself but is ‘acquired’ in the in vivo setting. This tuning can be parsimoniously explained by nasal odorant metabolism, with esters and aldehydes metabolized into their corresponding carboxylic acid by the enzymes carboxylesterase and aldehyde oxidase or aldehyde dehydrogenase. Alcohols may be metabolized into carboxylic acids via a two-step process, with alcohol dehydrogenase first generating the aldehyde followed by subsequent metabolism to the acid.

A strong prediction of the metabolism model is that acid-sensitive OSNs should be narrowly tuned to carboxylic acids within a narrow range of structural variation, but broadly tuned to different esters as long as the hydrolysis product of the ester yields the appropriate acid. The tuning of the 2-methyl-2-pentenoic acid-sensitive identified glomerulus to a range of tiglate esters, characterized earlier in (Burton et al., 2022), was consistent with this prediction (Fig. 2A). We tested the model further in awake mice (n = 6) by challenging the same glomerulus with an array of ester derivatives whose alkyl end spanned a broad range of structural features; we also included several aldehydes predicted to be metabolized into effective or ineffective carboxylic acids (Fig. 2B, C).

**Figure 2.**
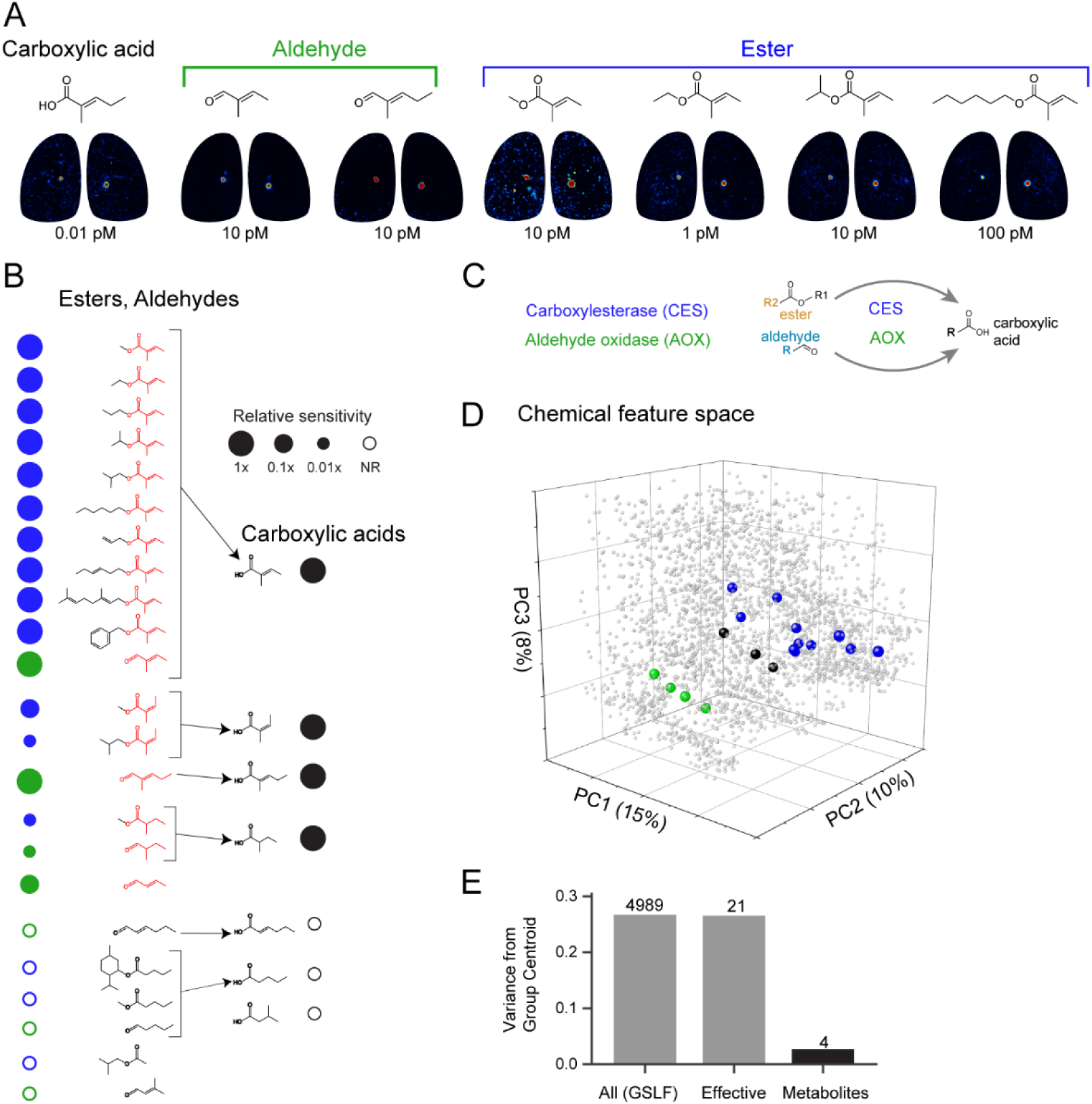
Odorant metabolism explains apparently broad tuning of OSNs across chemical feature space in vivo. **A.** OMP-GCaMP6s odorant response maps showing potent and selective activation of OSN inputs to a functionally-identified glomerulus with primary sensitivity to tiglic acid / 2-methyl-2-pentenoic acid and an array of esters with varying structural configurations of the alkyl end. Data from (Burton et al., 2022). **B.** Tuning of the same identified glomerulus to 30 odorants tested in awake mice. Sensitivity is expressed relative to the estimated reference concentration for tiglic acid of 0.1 nM (see Methods). Each odorant was screened using a serial 10-fold concentration series beginning at this estimated concentration; circles indicate lowest relative concentration eliciting a response. Open circles indicate no response at 100x reference concentration (10 nM). Colors indicate substrate for carboxylesterase or aldehyde oxidase/aldehyde dehydrogenase, respectively; arrows indicate carboxylic acid product. Data compiled from 6 mice (12 OBs): OMP-IRES-Cre x TIGRE2-GCaMP8m (n = 4);OMP-IRES-tTA x tetO-GCaMP6s (n = 2). **C.** Enzyme conversion of esters and aldehydes by carboxylesterase (CES) and aldehyde oxidase (AOX), respectively, each leading to production of the carboxylic acid derivative. Production of the alcohol by CES is not shown. **D.** Scatter plot of the 22 effective odorants from (B), in the chemical feature space defined by 4989 compounds (3637 shown,gray dots) taken from (Lee et al., 2023). Axes are principal components of MACCS chemical fingerprint features (PCs 1 - 3; 33% variance explained). Effective odorants are color coded by enzyme substrate or carboxylic acid product (black). **E.** Spread of effective odorants in PCs 1 - 4 of the MACCS feature space (39% total variance), compared to all GoodScents odorants and to only the metabolic carboxylic acid products. Effective odorants span nearly all of the chemical feature space, while their metabolic products span a much smaller portion.

We found that esters and aldehydes whose metabolic products matched effective carboxylic acids for the glomerulus were themselves effective odorants, while those predicted to be metabolized into ineffective acids were also ineffective (Fig. 2B). Collectively, the panel of effective odorants covered a large fraction of the ‘space’ of physicochemical odorant features, as defined by a database of 4989 odorous compounds (Lee et al., 2023) (Fig. 2D). However, all 22 of the effective esters and aldehydes are metabolized into one of four effective carboxylic acids, all of which activated the glomerulus. The four effective acids vary only slightly in structure and span a much smaller portion of chemical feature space (Fig. 2D, E). Thus, odorant metabolism contributes substantial explanatory power to accounting for the tuning of odorant receptors in physicochemical feature space.

### Odorant metabolism is rapid and predicts inhalation-linked timing differences in vivo

One reason for questioning the contribution of odorant metabolism to olfactory sensation is the presumption that this is a slow process relative to the dynamics of OSN activity relevant to perception, which occurs over the timescale of a single respiratory cycle and corresponds to a few hundred milliseconds for an awake mouse (Abraham et al., 2004; Spors et al., 2006; Schaefer and Margrie, 2007; Wesson et al., 2008; Shusterman et al., 2011; Wilson et al., 2017). To assess whether nasal metabolism can generate secondary odorants on this time-scale, we first measured the production of odorant metabolites directly from ex vivo olfactory epithelium, using Proton Transfer Reaction - Time-of-Flight Mass spectrometry (PTR-MS) measurements and a rapid odorant exchange device (Fig. 3A, Fig. S2A). The PTR-MS setup was optimized to detect the parent ester and its alcohol hydrolysis product; detection of the acid product was not feasible due to its low volatility.

**Figure 3.**
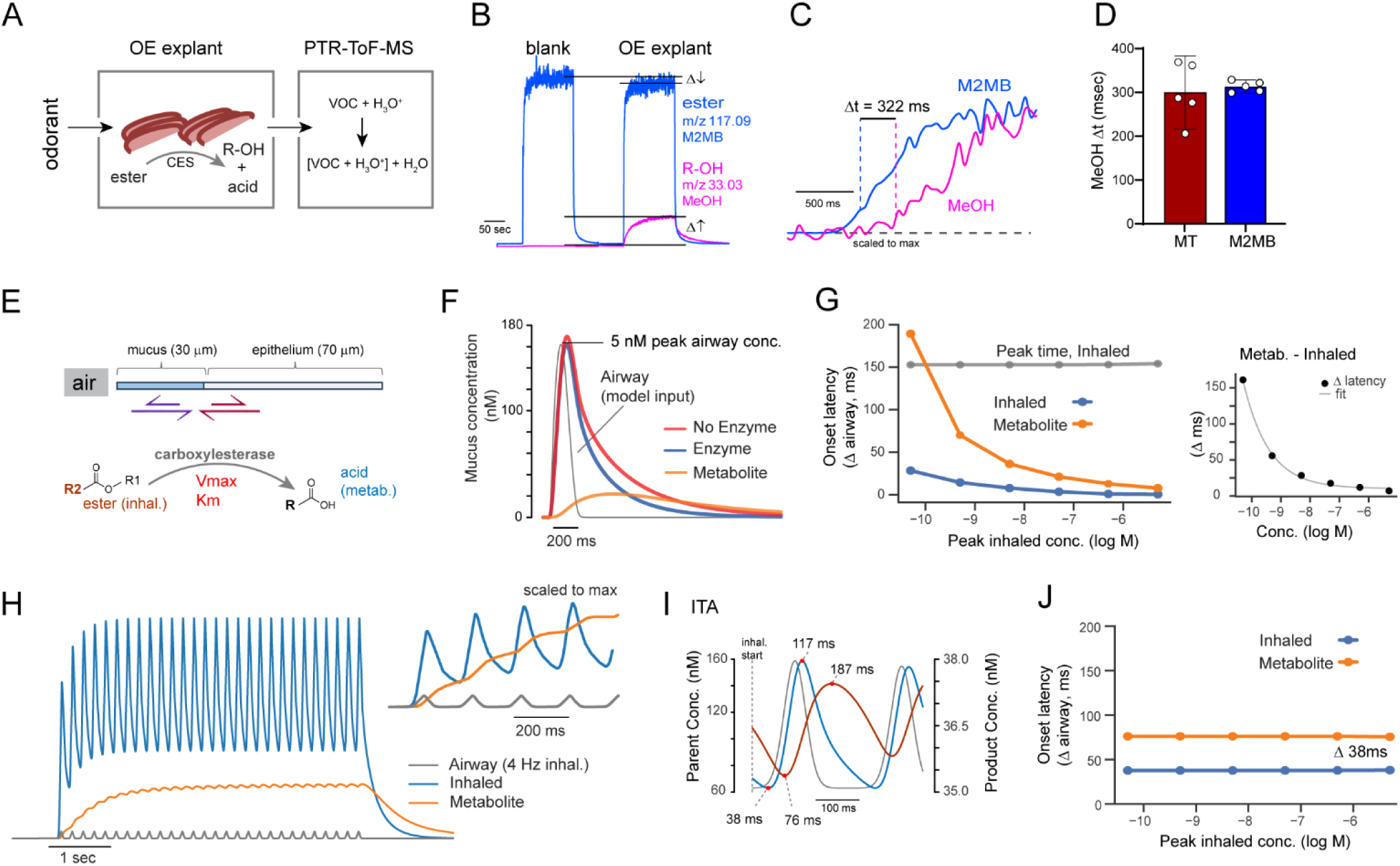
Nasal enzymes mediate rapid odorant metabolism on a time-scale relevant to inhalation-linked coding. **A.** Schematic of setup for direct measurement of ester metabolism by excised mouse nasal olfactory mucosa using PTR-MS. (VOC, Volatile Organic Compound). **B.** PTR-MS signal for the ester methyl-2-methyl butyrate (M2MB) and its methanol hydrolysis product, measured with and without olfactory mucosa present. In the presence of tissue, the methanol signal rapidly rises above control values within a few hundred milliseconds the 1-sec sampling period while the M2MB signal is reduced, consistent with ester metabolism. **C.** Expanded time-course of M2MB and methanol signal onsets from (B) and illustration of onset times and Δt. Each trace scaled to its own maximum. See Methods for details. **D.** Summary data for Δt measurements for methyl tiglate and M2MB. Bars indicate mean Δt; points are biological replicates; error bars show 95%CI. **E.** Schematic illustrating model for simulating absorption and metabolism of inhaled odorant during respiration in the mouse olfactory epithelium. See Text for details. **F.** Dynamics of inhaled odorant (methyl tiglate) concentration in the mucus compartment, with and without the presence of carboxylesterase enzyme, and metabolite product (methanol; orange), following a single inhalation. Dashed line shows odorant concentration profile in the airway, used as input to the simulation. Metabolite concentration rises with a delay of ∼ 100 ms and a slower decay. **G.** Onset latency for parent odorant and metabolite product as a function of peak inhaled concentration. Latency was defined as the time from inhaled odorant onset to a mucus concentration level of 1E-10 M. Gray trace shows time of peak mucus concentration for the inhaled odorant. Right: Parent-product latency difference versus concentration. Line shows fit to single-exponential decay, tau = 0.9 [log M], r^2^ = 0.998. **H.** Simulated odorant and metabolite dynamics during repeated inhalations at 4 Hz, showing accumulation of metabolite in the mucus over the first several inhalations, with weak modulation by each successive inhalation. **I.** Inhalation-triggered average (ITA) of inhaled odorant and metabolite during successive inhalations, showing delayed and less-modulated dynamics of the metabolite. **J.** Onset latency for parent odorant and metabolite product during successive inhalations, defined as trough of inhalation-triggered average (ITA) trace, as a function of peak inhaled concentration. In contrast to initial onset latencies, latencies for successive inhalations remain constant across concentration, with a Δt of 38 ms between parent and product.

PTR-MS sampling revealed production of the expected methanol hydrolysis product for methyl tiglate, methyl-2-methyl butyrate and methyl propionate, as well as a concomitant reduction in the concentration of the parent ester, relative to the no-tissue condition (Fig. 3B; Fig. S2C). We were not able to detect isopropanol from isopropyl tiglate and did not pursue this further. We focused on methyl tiglate and methyl-2-methyl butyrate for further analysis of the time-course of metabolite production. Using a scan frequency of 20 Hz (50 ms scan interval) and estimating the minimum delay (Δt) between the time of onset of the parent ester and detection of its methanol product (see Methods) yielded a Δt of 302 ± 30 ms (mean ± SEM.) for methyl tiglate (n = 5) and 313 ± 5 ms (n =5) for methyl-2-methyl butyrate (Fig. 3C, D). Given that a portion of Δt reflects setup-imposed delays including desorption of the alcohol from the mucosal explant and bulk flow through the chamber, these results indicate that mouse olfactory epithelium can generate appreciable metabolite odorants within the time-scale of a single respiratory cycle in an awake mouse breathing at 5 Hz.

To further estimate the time-course of metabolite odorant production on the time-scale of mouse respiration, we constructed a dynamic model that simulated the sorption of odorant from air into an aqueous mucus phase, its partitioning into the underlying epithelium, and odorant metabolism mediated by an enzyme (carboxylesterase) present in both compartments (Fig. 3E). The model used rate constants for ester metabolism experimentally determined in rodent nasal epithelium (Bogdanffy and Taylor, 1993), and input kinetics that mimicked odorant concentration dynamics during natural breathing (Jiang and Zhao, 2010). We focused on the model’s predictions of inhaled and metabolism-generated odorant concentrations in the mucus layer, which contains the OSN cilia and is the site of OR-ligand binding.

In response to a single inhalation of an ester odorant (methyl tiglate), concentration dynamics in the mucus followed that of the airborne odorant, albeit with a slightly delayed time to peak (Δt = 46 ms) and a slower decay (tau = 194 ms) (Fig. 3F). Increasing concentration of the inhaled odorant had minimal impact on the kinetics of the odorant in the mucus compartment and on a metric of latency defined as the time to reach an absolute threshold concentration (Fig. 3G). Introducing ester hydrolysis by carboxylesterase in the model minimally impacted ester concentration dynamics (decay tau = 175 ms) but produced a rise in mucosal concentrations of the acid metabolite that was delayed relative to the parent ester (Δ latency, 95 ms; Δ peak time, 346 ms) (Fig. 3F). Concentration substantially impacted response latency of the metabolite odorant, reflecting the concentration-dependence of the rate of odorant metabolism and the accumulation of metabolite in the mucus, with the latency difference between the inhaled and metabolite odorants well-fit by an exponentially decaying function (Fig. 3G).

Simulating odorant concentration profiles during repeated inhalations at 4 Hz revealed an accumulation of metabolite over the first few inhalations (Fig. 3H). Metabolite concentration was still modulated by each inhalation, but the degree of modulation was much weaker and lagged behind that of the inhaled odorant (Fig. 3H). The lag between the inhaled odorant and its metabolite ranged from ∼40 - 70 ms (lag of cross-correlation between inhalation-triggered average waveforms, 60 ms) (Fig. 3J).

These simulations, while simplified, nevertheless indicate that nasal metabolism can generate secondary odorants with sufficient speed to activate metabolite-sensitive OSNs within the time-scale of a single respiratory cycle, but with delayed and dampened respiratory patterning in metabolite concentration across successive inhalations. They also suggest that the impact of concentration on the latency of responses to inhaled odorants is quite limited, with a dynamic range well under 50 ms.

To test predictions of the metabolism model in vivo, we analyzed OSN response dynamics relative to inhalation timing in the awake mouse. We monitored respiration via an external sensor and presented odorants using a custom olfactometer that achieved rapid and reliable onsets in the interval between inhalations (Supplementary Figure S3). After aligning OMP-GCaMP8m signals to the first inhalation after odorant onset, latencies measured from the four functionally-identified glomeruli (at 10 - 100x threshold concentration) fell within a narrow range for their respective ‘primary’ carboxylic acids (TA, 2-MBA, PA, IVA; mean latencies (n replicates): 103 (6), 77 (6), 89 (3) and 93 (4) ms, respectively) and were significantly longer for the ester, aldehyde and alcohol derivatives (Fig. 4A-C; Suppl. Fig. S4), with delays relative to the acid of 98 ± 11 ms (*p* = 2E-7); 88 ± 11 ms (*p* = 1.5E-6) and 205 ± 38 ms (*p*=7.0E-4), respectively (estimated marginal means ± SEM, mixed effects model; details in Methods).

**Figure 4.**
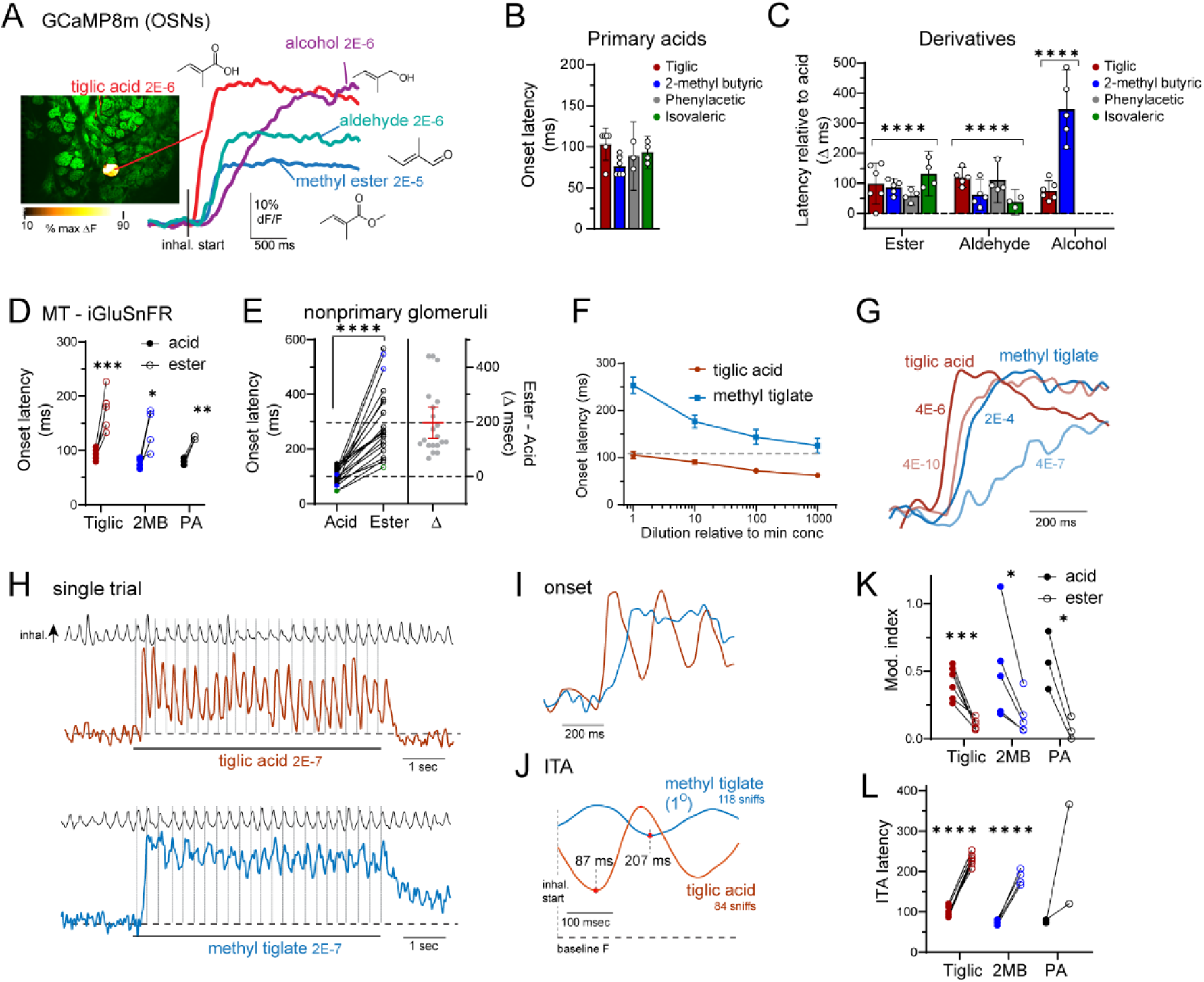
Inhalation-linked timing of OSN activation by acids and their derivatives in the awake mouse is consistent with the odorant metabolism model. **A.** OSN response onsets (OMP:GCaMP8m) to tiglic acid and its ester, aldehyde and alcohol derivatives, aligned to the first inhalation of each odorant. Responses to the derivatives are delayed relative to the acid, with tiglic alcohol showing the largest delay. Same preparation as in Fig. 1C. Numbers indicate total odorant dilution. **B.** Summary data showing similar onset latencies in each of the four functionally-identified glomeruli to their respective primary carboxylic acid odorant. **C.** Summary data showing delays in onset latency for the ester, aldehyde and alcohol derivatives for the four glomeruli, relative to that of the acid measured in the same session. Data presented as mean ± 95% CI; points indicate biological replicates; n = 3 - 6 replicates per glomerulus. Linear mixed-effects model followed by Bonferroni-corrected pairwise comparisons of estimated marginal means (EMMs) between acids and their corresponding esters, aldehydes, and alcohols. \*\*\*\**p* < 0.0001. **D.** Summary of MT-iGluSnFR onset latencies in responses to each primary acid odorant and its ester derivative for three of the functionally-identified glomeruli. Data show paired measurements in biological replicates. Asterisks, paired t-test results per odorant pair; \*\*\**p* = 0.001; \*\**p*<0.01; \**p*<0.05. **E.** Paired comparison of onset latencies for each carboxylic acid and its ester derivative, measured in ‘nonprimary’ glomeruli recruited at suprathreshold concentrations. All acid-sensitive glomeruli respond with longer latencies to their ester derivative. Right plot shows estimation plot of differences, error bars, 95%CI. Linear mixed-effects model followed by pairwise comparison between acids and esters using estimated marginal means; n = 20 glomeruli. \*\*\**p* < 0.001. **F.** iGluSnFR3 onset latencies for tiglic acid and its ester derivative as a function of concentration, measured from replicates of the tiglic acid-sensitive identified glomerulus. Dashed line, latency for tiglic acid at near-threshold concentration, for reference. Points and error bars indicate mean ± SEM. **G.** Timing of sensory input to mitral/tufted (MT) cells imaged with iGluSnFR3 in functionally-identified glomeruli. Traces show inhalation-aligned responses to tiglic acid (TA, red) and methyl tiglate (MT, blue) presented at near-threshold and suprathreshold (1000 - 10000x) concentrations. Note that MT responses are slower than TA responses at all concentrations. **H.** Single-trial traces of iGluSnFR3 signals during presentation of tiglic acid and methyl tiglate in the awake mouse, taken from primary glomerulus. Top trace shows respiration signal; vertical lines indicate start of each inhalation during odorant presentation. **I.** Expansion of response to odorant onset, from trial in (H). Acid-evoked signals are strongly modulated by each inhalation, while ester signals are slower-onset and show weaker modulation. **J.** Inhalation-triggered average (ITA) traces of iGluSnFR3 signals during successive inhalations of tiglic acid and methyl tiglate, showing delayed inhalation-linked onset (defined by trough of ITA) and weaker modulation for the ester. ITA traces generated from same session as in (H). Note that MT ITA response is delayed by greater than one respiratory cycle (dashed line at 240 ms), such that an early trough is apparent reflecting the response to the preceding inhalation (dashed line at 27 ms). **K.** Paired comparison of modulation index metrics measured from ITA traces for the acid and ester derivatives, plotted for each primary odorant as in (D). Points indicate biological replicates. Asterisks, paired t-test per odorant pair: \*\*\**p* = 0.002; \**p*<0.05. **L.** Paired comparison of ITA latencies for the primary acid and ester derivative for the three functionally-identified glomeruli, plotted for each primary odorant as in (D). Paired t-tests per odorant pair: **** p < 1e-4.

To characterize the inhalation-linked dynamics of sensory input with higher fidelity than reported by GCaMP8m, we used the genetically-encoded glutamate sensor iGluSnFR (A148S or iGluSnFR3 variants) (Marvin et al., 2018; Aggarwal et al., 2023), expressed via viral vector in mitral and tufted (MT) cells and imaged from their dendritic tufts (Moran et al., 2021b). iGluSnFR is optimal for its fast decay rate and high dynamic range and ability to follow sensory-driven transients at frequencies corresponding to natural breathing in awake mice (Aggarwal et al., 2023). MT- iGluSnFR signals predominately reflect OSN-driven glutamate signaling, with a lesser contribution from external tufted cells (Moran et al., 2021a).

Consistent with the presynaptic data, odorant-evoked glutamate onsets in functionally-identified glomeruli were slower for the ester derivative than for the primary carboxylic acid (mean Δacid 141 ± 13 ms (emm ± SEM), n = 14; *p* = 3E-6) (Fig. 4D, E). Aldehyde derivatives, tested in a few preparations, also evoked slower responses (tiglic aldehyde, Δacid = 33 ms, n = 2; phenylacetaldehyde, Δacid = 206 ms, n = 1). Acid-ester co-tuning and latency differences generalized beyond the most-sensitive glomeruli: 80% of glomeruli recruited by suprathreshold concentrations of a carboxylic acid (’nonprimary glomeruli’) were also activated by its ester derivative (MT-iGluSnFR: 20/25 glomeruli, 3 mice, 3 acid-ester pairs;OMP-GCaMP8m: 48/60 glomeruli, 7 mice (8 OBs), 4 acid-ester pairs); the remaining 20% may reflect glomeruli with low sensitivity to the acid such that the ester does not produce sufficient acid metabolite at the concentration presented. Notably, all acid-ester co-tuned glomeruli showed longer-latency responses to the ester (MT-iGluSnFR: Δacid = 197 ± 27 ms (mean ± SEM), *p* < 1E-6; Fig. 4E; OMP-GCaMP8m: Δacid = 143 ± 22 ms, *p*< 1E-6 ms; Fig. S4J).

Longer latencies might be explained by a difference in sensitivity to the acid versus its derivatives. Thus we tested each primary acid-ester pair at dilutions ranging from near-threshold for signal detection to 2 – 3 log units more concentrated. Acid-evoked onset latencies decreased modestly with increases in odorant concentration (Fig. 4F, Suppl. Fig. S4), decreasing from 10 - 30 ms per ten-fold concentration increase (median, 21 ms). Two different functionally-identified glomeruli with primary sensitivities to ketones (2-hexanone and 2’-OH acetophenone, respectively) showed a similar latency decrease across concentration (Suppl. Fig S5). Onset latencies to the ester derivative decreased with increasing concentration but remained 50 - 100 ms longer than acid-evoked latencies at a given relative concentration (Fig. 4F; Suppl. Figure S4). In some cases, responses to the highest concentrations of methyl tiglate (up to 1000x perithreshold) were slower than those evoked by near-threshold concentrations of tiglic acid (Fig. 4G). GCaMP8m signals in OSN terminals showed a similarly modest concentration-dependence, (Figure S4), save for a shift to longer latencies at near-threshold concentrations likely reflecting summation across multiple inhalations captured by the slower decay of GCaMP8m (Zhang and Looger, 2024).

Across successive inhalations of odorant, MT-iGluSnFR signals evoked by carboxylic acids showed prominent transients time-locked to each inhalation, while responses in the same glomerulus to the parent ester showed much weaker inhalation-linked modulation (Fig. 4H, I). To quantitatively compare inhalation-linked patterning, we generated inhalation-triggered average (ITA) response waveforms by averaging the MT-iGLuSnFR in a 400 ms snippet following each inhalation after the first 500 ms of odorant presentation (Fig. 4J). We excluded any inhalations occurring at a frequency above 5 Hz, yielding 50 - 150 inhalations over 2 - 6 repeated trials. The trough of the ITA waveform provided a robust measure of onset latency for repeated inhalations of odorant. To quantify the degree of inhalation-linked modulation independent of latency, we defined a modulation index (MI) metric with an MI of 1 reflecting responses that return completely to baseline with each inhalation and an MI of 0 representing tonic responses with no inhalation-linked patterning (Methods).

In all three identified glomeruli, inhalation-linked modulation of responses to the ester was significantly weaker than for the carboxylic acid and sometimes absent altogether (i.e., MI ∼ 0) (Fig. 4J, K) (Tiglic, *p* = 0.003, n = 6; 2-methyl butyric, *p* = 0.034, n = 5; Phenylacetic, -0.5 ± 0.07, *p* = 0.022, n= 3; paired t-tests; mean Δ MI across group (emm ± SEM), -0.36 ± 0.05, n = 14, *p* = 5.5E-6, Linear mixed-effects model with pairwise emmeans comparison). ITA response latencies (when measurable modulation was present) were also longer for the ester compared to the acid (Fig. 4L; Tiglic, *p* = 2E-6; n=6; 2-methyl butyric, *p* = 7E-5; n=5; mean Δ latency across group (emm ± SEM), 128 ± 15 ms, n = 13; *p* = 1.6 E-6, Linear mixed-effects model with pairwise emmeans comparison). In some cases, latencies exceeded the duration of the average respiratory cycle (i.e., 200 ms at 5 Hz respiration), resulting in inhalation-triggered latencies that were paradoxically shorter for the ester than the acid (see Fig. 5D), and less than 30 ms after inhalation onset, which is difficult to reconcile with odorant transduction and axonal conduction delays (Wesson et al., 2008).

**Figure 5.**
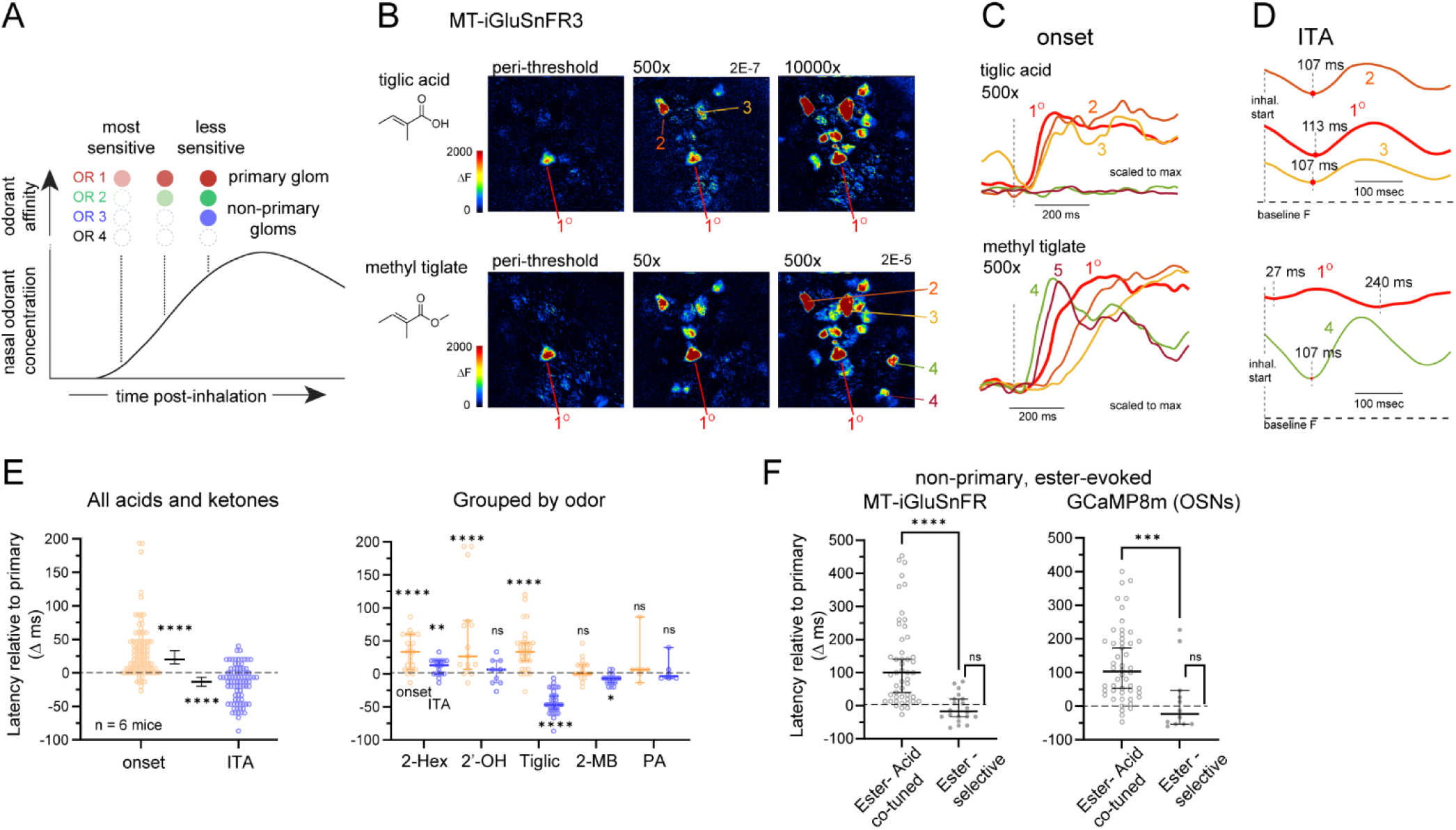
Odorant metabolism alters sensitivity-timing relationships among OSN inputs to OB glomeruli. **A.** Schematic of sensitivity-latency model determining inhalation-linked timing of OSN inputs as a function of odorant concentration and relative sensitivity, tested in the subsequent figure panels. **B.** MT-iGLuSnFR3 response maps showing primary (i.e., most-sensitive) glomerulus for tiglic acid (TA) and its ester derivative methyl tiglate (MT) and recruitment of non-primary glomeruli at higher odorant concentrations. **C.** Overlay of inhalation-aligned response onsets in primary and non-primary (less-sensitive) glomeruli evoked by 500x threshold concentration of TA or MT. Top: Non-primary responses to TA are slightly delayed relative to the primary glomerulus. Bottom: Two non-primary glomeruli (ROIs 4 and 5) respond with shorter latency to the primary glomerulus, despite being ∼500x less sensitive. **D.** ITA traces generated from the same trials as in (C). Top: Primary and non-primary glomeruli show near-identical latencies during successive inhalations of TA. Bottom: The shorter response latency in the non-primary ROI 4 (from C) persists during successive inhalations and is identical to that of TA-evoked ITA responses (compare to top traces). **E.** Odorant onset and ITA latencies for all non-primary glomeruli recruited by suprathreshold concentrations of carboxylic acids or ketones, relative to the primary glomerulus. Left: Across all odorants, onset latencies are slightly longer, while ITA latencies are slightly but significantly shorter than their respective primary glomeruli. Right: Same data grouped by odorant shows variability in the magnitude of sensitivity-timing relationships for different odorants. Right: Same data, grouped by odorant and showing statistical comparison to zero (same latency as primary glom.). ****p < 1e-4, ***p < .001, **p < .01, *p<.05, Emmeans post-hoc test, Tukey adjustment for multiple comparisons. **F.** Latency-sensitivity relationships differ substantially for ester-responsive non-primary glomeruli depending on whether they are co-tuned to their acid hydrolysis product (’Ester co-tuned’) or not (’Ester-selective’). \*\*\*\**p* < 1E-4; \*\*\**p* < 1E-3, Mann-Whitney test.

Taken together, these results support a model in which nasal odorant metabolism rapidly generates secondary odorants that robustly activate OSNs on a time-scale relevant to timing-based odor coding strategies.

### Odorant metabolism alters sensitivity-timing relationships among OSN inputs in vivo

Variance in inhalation-linked response latencies has been proposed to reflect the relative sensitivity of a receptor (or its cognate glomerulus) to a given odorant (Wilson et al., 2017). A core prediction of this ‘sensitivity-latency’ model is that glomeruli receiving the most-sensitive, ‘primary’ OSN inputs remain activated earliest across concentration, with lower-sensitivity glomeruli responding later than their higher-sensitivity counterparts (Fig. 5A), enabling a timing-based strategy for concentration-invariant odor identity coding (Hopfield, 1995; Wilson et al., 2017). To test how odorant metabolism may impact such a strategy, we analyzed sensitivity-timing relationships across glomeruli for each carboxylic acid and its metabolic parent ester by varying odorant dilutions from near-threshold for signal detection to 2 – 3 log units more concentrated. We included two additional functionally-identified glomeruli with primary sensitivities to non-acid (ketone) odorants (2-hexanone, and 2’-OH acetophenone) (Burton et al., 2022). Estimated delivered perithreshold concentrations ranged from 1E-14 M to 1E-9 M (Table S1).

Figures 5B-D show example response maps and inhalation-aligned MT-iGLuSnFR traces from primary and non-primary glomeruli activated by tiglic acid and its parent ester, methyl tiglate. Near-threshold concentrations of either odorant evoked input to the same single glomerulus in the field of view, identifying it as the ‘primary’ glomerulus, while increasing concentrations recruited input to additional, ‘non-primary’ glomeruli (Fig. 5B). Similar concentration-dependent recruitment was seen for the other functionally-identified glomeruli (Figure S5).

We compared onset latencies of less-sensitive, ‘non-primary’ glomeruli with that of the ‘primary’ glomerulus, imaged in the same suprathreshold concentration trials (Fig. 5C, D, E). In response to carboxylic acids and ketones, non-primary onset latencies were, overall, longer than those of the primary glomerulus - consistent with the sensitivity-latency model - but latency differences were small (Fig. 5C, top traces; Fig. 5E): the median delay across all non-primary glomeruli, relative to the primary glomerulus imaged in the same field of view, was 20 ms (IQR: 7 - 47 ms; *p* < 1E-6; one-sample Wilcoxon Signed Ranks, n = 96 glomeruli, 7 mice, 8 OBs), with no significant delay for some odorants (e.g., Phenylacetic acid, Fig. 5E, right). Latency differences skewed longer for OSN-GCaMP8m signals, with a median non-primary delay of 47 ms (IQR: 13 - 80 ms; *p* < 1E-6; one-sample Wilcoxon Signed Ranks, n = 56 glomeruli, 6 OBs, 5 mice)).

The sensitivity-latency relationships apparent at odorant onset did not persist across subsequent inhalations. In fact, across all non-primary glomeruli, inhalation-triggered latencies beyond the first 500 ms of odorant presentation were significantly *shorter* than those of the primary glomerulus (median ITA latency relative to primary: -13 ms; IQR: -40 - 3 ms; *p* = 4E-6; one-sample Wilcoxon Signed Ranks test, n = 85 glomeruli, 7 mice, 8 OBs) (Fig.5D, E). The change in sensitivity-based timing differences after the first inhalation differed for different odorants (Fig. 5E) (One-way ANOVA, onset latency: *p* < 2E-16, F(4,80) = 47.7; ITA latency: *p* = 0.002; F(4,91) = 4.5). Tiglic acid and 2-methyl butyric acid showed the most consistent reversals in relative latency values, with median delays of -47 and -7 ms, respectively, *p* < 1E-4; one-sample Wilcoxon Signed Ranks). The loss of timing differences between primary and non-primary glomeruli may reflect adaptation of the most-sensitive glomerulus to suprathreshold odorant concentrations. Overall these results suggest that latency does not robustly distinguish the most-sensitive from less-sensitive glomeruli across repeated inhalations of odorant.

We asked how odorant metabolism impacts sensitivity-timing relationships by comparing primary and non-primary latencies among glomeruli activated by the parent ester for each of the three carboxylic acids. At suprathreshold concentrations, non-primary glomeruli recruited by the ester showed a larger range of latency differences (relative to the primary glomerulus) than the same non-primary glomeruli recruited by the acid (Fig. 5F). At the same time, a few of the non-primary glomeruli recruited by the ester responded at short latency. Notably, these glomeruli were distinguished by being *insensitive* to the corresponding acid, and thus were labeled ‘ester-selective’. Latencies for ester-selective nonprimary glomeruli were not significantly different from that of the primary glomerulus (MT-iGLuSnFR: *p* = 0.34, n = 20; OMP-GCaMP8m: *p* = 0.73, n = 12; one-sample Wilcoxon Signed Ranks) and were significantly shorter than glomeruli co-tuned to the ester and its carboxylic acid derivative, imaged in the same trials (Fig. 5F).

Many ester-selective glomeruli responded with latencies *shorter* than that of the primary glomerulus; examples of two such glomeruli recruited by methyl tiglate are shown in Fig. 5B and C (ROIs 4 and 5). Such glomeruli were 100 - 1000x less sensitive to the ester than the primary glomerulus, yet responded up to 60 ms *earlier* (Fig. 5F), representing violations of the sensitivity-latency model. These timing differences persisted for subsequent inhalations of the ester and were associated with strong inhalation-linked modulation (Fig.5D). Similar results were seen for ester-evoked responses in OMP-GCaMP8m signals (Fig. 5F).

### Inhalation-linked timing disambiguates glomerular representations of inhaled versus metabolism-generated odorants

To assess the relationship between chemical tuning and response timing across a larger array of glomeruli, we returned to wide-field imaging of OSN-GCaMP8m signals across the full extent of the dorsal OB, mapping responses to acids and their ester derivatives. At suprathreshold concentrations, the acid and ester activate nearly the same set of glomeruli in the anterior-medial OB, consistent with the co-tuning already described; esters also activated glomeruli in the caudal-lateral OB that were not activated by the acid (Fig. 6A). These clusters of distinct tuning were contained within the respective domains of class I or class II OR-expressing OSN projections, which we defined functionally using high concentrations of a carboxylic acid (butyric acid) or ketone (2-hexanone), as done previously (Bozza et al., 2009) (Fig. 6B).

**Figure 6.**
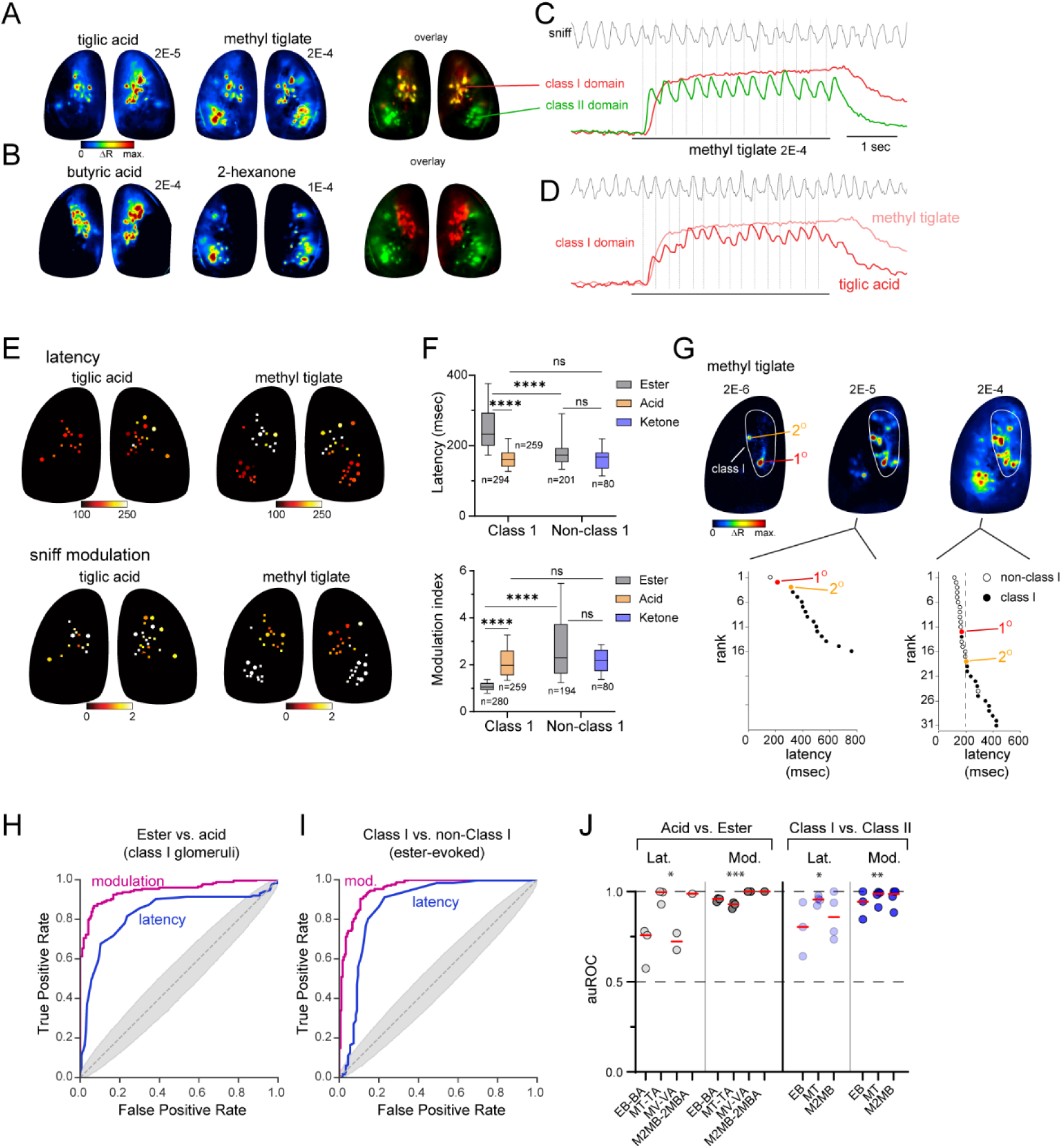
Inhalation-linked timing maps to OR-defined topographic domains and disambiguates glomerular representations of inhaled versus metabolism-generated odorants. **A.** Widefield OSN-GCaMP8m response maps to suprathreshold concentrations of the acid TA and its ester derivative MT. Red-green overlay (right) shows near-complete overlap of TA and MT - responsive glomeruli in and anterior-medial OB and selective activation of caudal-lateral glomeruli by MT. **B.** Response maps from the same mouse to suprathreshold concentrations of a carboxylic acid (butyric acid) and ketone (2-hexanone). Red-green overlay illustrates nonoverlapping glomerular activation defining class I and class II-expressing OSN projection domains. **C.** GCaMP8m traces imaged from one glomerulus in the class I (red) and class II (green) domains, in response to MT in the same trial. Top trace shows mouse respiration, vertical lines indicate beginning of each inhalation. Note slower onset and lack of modulation in the class I - located glomerulus. **D.** Single-trial trace from the same class I - located glomerulus in (C) to TA (red trace), showing fast onset and faithful modulation by each inhalation. Response to MT (from C), aligned to the first inhalation of odorant for both, is shown for comparison. **E.** Maps of glomeruli evoked by TA or MT (same data as in A), pseudocolored by onset latency (top) or modulation index (bottom), showing topography of each metric matching OR class-defined domains. **F.** Comparison of onset latency (top) or modulation index (bottom) for ester or acid-evoked responses, grouped by glomerular location (class I domain or non-class I location). Asterisks indicate unpaired t-test results, \*\*\*\**p* < 1E-4.: class I acid vs. ester latencies: 177 ± 4 ms vs. 260 ± 6 ms, p < 1e-6; class I vs. non-class I ester latencies: 260 ± 6 (SEM.) ms vs. 199 ± 8 ms, p < 1e-4. acids vs. 2-hexanone: 177 ± 4 ms vs. 173 ± 8 ms, p = 0.71; class I acid vs. ester mods: 2.15 ± 0.5 vs. 1.07 ± 0.02, p < 1e-4; class I vs. non-class I ester mods: 1.07 ± 0.02 vs. 2.9 ± 0.14, p < 1e-4; acids vs. 2-hexanone mods: 2.13 ± 0.05 vs. 2.2 ± 0.07, p = 0.55. Acids tested: tiglic acid, 2-methyl butyric acid, butyric acid, valeric acid; esters: methyl tiglate, methyl-2-methyl butyrate, ethyl butyrate, methyl valerate. **G.** Top: Widefield response maps to three concentrations of MT (100-fold range) reveal widespread violations of sensitivity-timing relationships. The glomeruli most sensitive to MT (1°, 2°) are located within the class I domain. Bottom: Dot plots showing ranked latencies of glomeruli activated by suprathreshold concentrations of MT, with 1° and 2° glomeruli highlighted. Filled or open dots indicate glomeruli in the class I or class II domains. **H.** Receiver-operating characteristic (ROC) curves showing performance for discriminating activation of a glomerulus by a carboxylic acid or its ester derivative on the basis of onset latency or modulation index. Shaded area indicates envelope of ROC curves for scrambled data. n = 346 glomeruli from 6 OBs (4 mice). **I.** Same analysis for classifying glomerular identity as class I (acid-sensitive) or class II (ester-selective) for ester-responsive glomeruli on the basis of latency or modulation index. n = 415 glomeruli from 6 OBs (4 mice). **J.** Performance scores (auROC) for specific acid-ester pairs (left) or specific esters (right). Points indicate auROC calculated from one OB. Red line, median. Asterisks indicate significant classification (auROC > 0.5) across all acid-ester replicates within a group. \*\*\**p* < 1e-3, \*\**p* <.01, \**p*<0.05. Linear mixed-effects model with one-sided emms post hoc test (null = 0.5 auROC).

Mapping onset latency or sniff-modulation (MI) to glomerular location revealed a topography of inhalation-linked timing features corresponding to each domain. As seen in the example for suprathreshold tiglic acid and methyl tiglate (Fig. 6C, D, E), glomeruli in the caudal-lateral class II domain responded to suprathreshold ester concentrations with short latency and strong modulation by each inhalation, while class I domain glomeruli responded with longer latency and little to no modulation in the same trials. In response to carboxylic acids (e.g., tiglic acid, Fig. 6D, E), class I glomeruli were activated with short latency and stronger modulation. This result was observed across four different acids and their ester derivatives (Fig. 6F, see legend for statistical test results; 7 OBs in 4 mice). We also mapped latency and sniff-modulation (MI) for the class II-selective ketone 2-hexanone; latencies and MI for 2-hexanone responses were not significantly different from those of class I – domain responses to acids and class II-domain responses to esters (Fig. 6F), indicating that inhalation-linked dynamics are not determined by glomerular location, OSN class or odorant class alone. These results are explained by rapid ester metabolism via nasal carboxylesterase, in which the carboxylic acid product of ester hydrolysis activates class I OSNs with a modest delay and dampened modulation, while class II OSNs responding directly to the ester respond with short latency and strong modulation with each inhalation.

Given that only a fraction of inhaled ester is hydrolyzed to produce the carboxylic acid product, one might predict that ester-evoked responses in class I glomeruli would only occur at high concentrations - higher, at least, than threshold concentrations for glomeruli sensitive to the ester itself. Surprisingly, however, we found that esters activated class I - domain glomeruli at lower concentrations than class II glomeruli (Fig. 6G) - at least those visible on the dorsal OB. As a result, recruitment of class II glomeruli at suprathreshold concentrations elicited many violations of the sensitivity-timing model. For example, in response to a ∼100x suprathreshold concentration of methyl tiglate, approximately half of the responsive glomeruli showed latencies *shorter* than those of the two most-sensitive (i.e., primary) glomeruli (Fig. 6G) (rank percentile: 48 ± 4 %-ile (mean ± SD), raw rank: 12 ± 2 of 24 ± 5 ROIs; n = 7 OBs, 4 mice). One explanation for these results may be extreme sensitivity of class I ORs to their carboxylic acid ligands (see Discussion).

Taken together, these results suggest that nasal odorant metabolism generates confounds for receptor-determined models of odor coding by distorting sensitivity-timing relationships and generating similar patterns of OSN activation for odorants with distinct structures. To evaluate the degree to which inhalation-linked timing can resolve these confounds, we quantified the ability of onset latency or sniff-modulation (MI) to classify an odor source as directly-inhaled or metabolically-generated, using receiver-operating characteristic (ROC) analysis. We first analyzed glomeruli co-activated by a given carboxylic acid and its metabolic parent ester (all presumed class I-OSN glomeruli), taken from widefield suprathreshold responses to each. Both onset latency and MI discriminated acids from their ester derivatives well above chance (Fig. 6H, J, Supp. Fig. S6; paired acids vs. esters: latency, *p* = 0.02; MI, *p* = 1.4E-3), and were able to classify ester-evoked responses as arising either from class I– or class II– domain glomeruli (reflecting metabolism- or directly-inhaled responses, respectively) (Fig. 6I, J Suppl. Fig. S6; Class I vs. Class II: latency, *p* = 0.015; MI, *p* = 1.6E-3). For some acid-ester pairs, the distribution of latency or MI values, taken from the same glomeruli in the same session, was nonoverlapping (auROC = 1; perfect classification). Thus, inhalation-linked timing can robustly discriminate neural responses to inhaled odorants from those of their metabolites.

### Neural representations of metabolism-generated odorants persist at the level of OB output

Finally, we asked whether neural circuits in the OB might differentially process inhaled versus metabolism-generated afferent signals on the basis of their on inhalation-linked timing. This prediction arises from a recent report of OB suppression of delayed OSN inputs on a time-scale matching the delay between inhaled and metabolism-generated odorants (Karadas et al., 2026). To test this prediction, we returned to the functionally-identified glomeruli, probing their mitral and tufted cell (MT) outputs using GCaMP8f, expressed in MT cells via viral vector in Tbet-Cre mice. Glomeruli were identified using the same diagnostic, low-concentration ‘primary’ odorants as for OSN imaging while imaging GCaMP signals from MT dendritic tufts in the glomerular layer (Fig. 7A).

**Figure 7.**
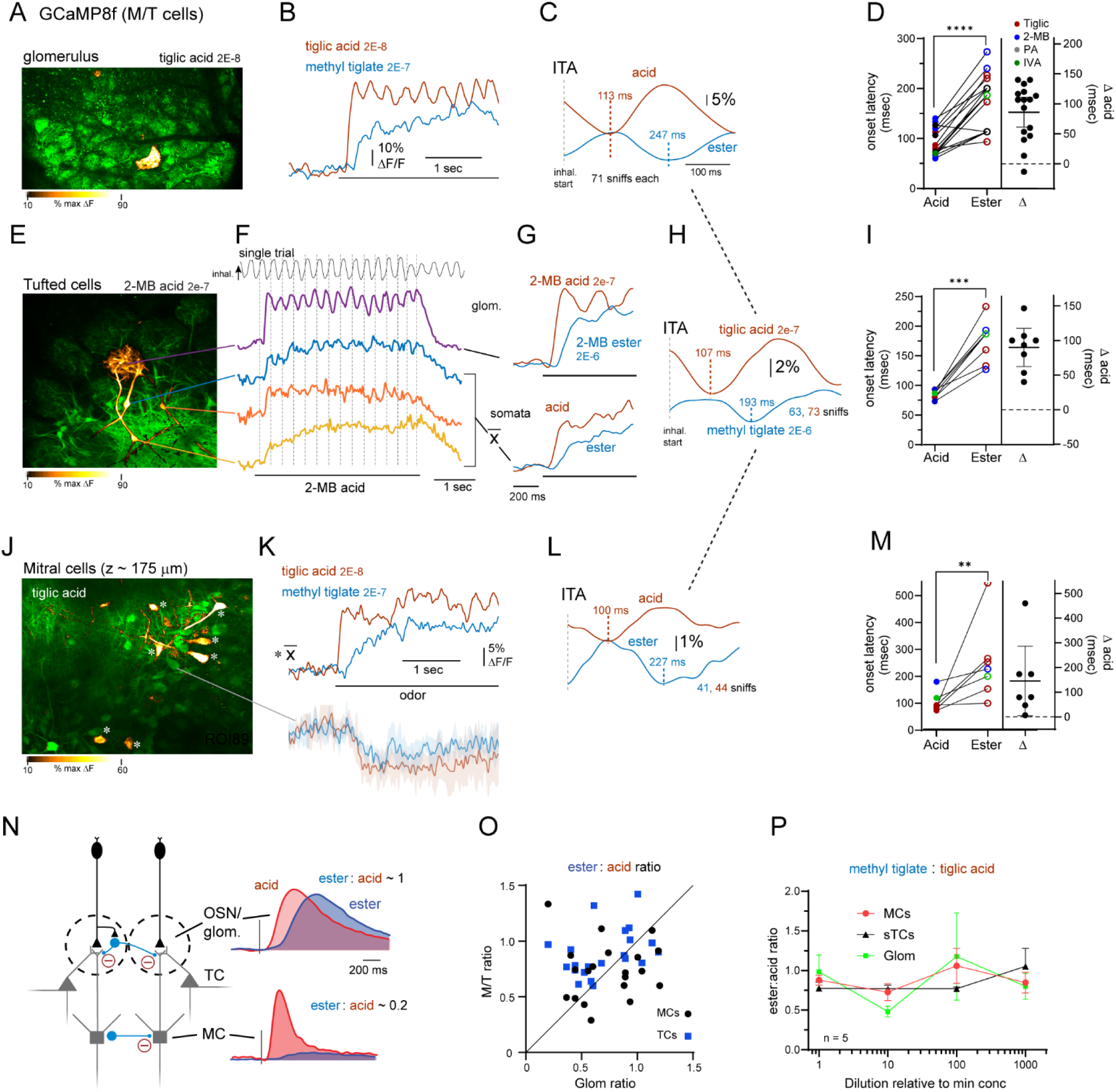
Delayed sensory inputs persist as delayed outputs among OB mitral and tufted cells. **A.** Response map (ΔF) of GCaMP8f signal in M/T cell dendritic tufts in a single glomerulus evoked by near-threshold concentration of tiglic acid, characteristic of this functionally-identified glomerulus identified glomerulus. **B.** Inhalation-aligned response onsets to TA and MT (10x threshold concentration), showing delayed response to the ester, as described for presynaptic signals. **C.** Inhalation-triggered average trace for GCaMP8f response to successive inhalations of TA and MT, with a delayed inhalation-linked modulation onset to the ester. **D.** Paired latencies measurements and difference estimation plot (mean ± 95%CI) for the primary acid and its ester derivative. Points of same color are biological replicates per glomerulus-primary odorant pair. ****, *p* < 1e-4; Mixed-effects model, post-hoc pairwise emms comparison, n = 16. Δ acid (emm ± SEM) = 87 ± 12 ms. **E.** Response map of GCaMP8f signal evoked by near-threshold 2-methyl butyric (2-MB) acid, showing singular activation of this identified glomerulus and simultaneous measurement of GCaMP8f signal in the glomerular tuft and three daughter tufted cells (TCs). **F.** Time-course of signal in the glomerulus and each daughter TC, imaged in the same trial. Top trace and dashed lines show respiration measurement and inhalation onset times. Note lack of clear modulation by inhalation in somata as compared to dendritic tufts. **G.** Inhalation-aligned response onsets to 2-MB acid and its methyl ester (2-MB ester) for GCaMP8f signals in the tuft (top) and averaged across the three daughter TCs (bottom), showing slower ester-evoked responses in both compartments. **H.** ITA trace for GCaMP8f response to successive inhalations of tiglic acid and methyl tiglate (same preparation as (A) and (J)), showing delayed inhalation-linked modulation onset to the ester. ITA waveform taken from mean signal across 9 daughter TCs from glomerulus in (A) and indicated number of inhalations. **I.** Paired latency measurements and difference estimation plot for the primary acid and its ester derivative in cohorts of daughter TC somata innervating identified glomeruli, plotted as in (D). *p = 0.001; Mixed-effects model, post-hoc pairwise emms comparison, n = 8 parent glomeruli. Δ acid (emm ± SEM) = 90 ± 12 ms. **J.** Response map of GCaMP8f signal in daughter mitral cells (MCs) of the same tiglic acid-identified glomerulus shown in (A). **K.** Top: Inhalation-aligned response onsets to TA and the MT ester (10x threshold concentration), averaged across 7 daughter MCs, showing delayed excitation by the ester. Bottom traces show GCaMP8f signal in an adjacent MC that is suppressed by both odorants. Shaded area, 95% CI across trials. **L.** Inhalation-triggered average trace for GCaMP8f response to successive inhalations of TA and MT (from same trials as in (K)), with delayed inhalation-linked modulation onset to the ester. **M.** Paired latency measurements and difference estimation plot for the primary acid and its ester derivative in cohorts of daughter MC somata innervating identified glomeruli, plotted as in (D). ***p = 0.044; Mixed-effects model, post-hoc pairwise emms comparison, n = 7 parent glomeruli. Δ acid (emm ± SEM) = 146 ± 57 ms. **N.** Schematic illustrating OB inhibitory circuits and predicted suppression of delayed sensory inputs, along with analysis metric for comparing net excitation ratios for acid versus ester responses. **O.** Scatter plot of Ester : Acid excitation ratios for intensity-matched ester - acid pairs measured from the glomerular tuft (X-axis, ‘Glom ratio’) versus the cohort of daughter TCs or MCs measured in the same experiment (Y-axis, ‘M/T ratio’). MCs show no change in excitation ratio relative to their parent glomerular signal, while TCs show a slight increase, indicating slightly stronger excitation in TC responses to the ester. **, *p* < 0.01; n.s., not significant, Wilcoxon Signed Ranks. **P.** Ester : Acid excitation ratio as a function of odorant intensity (matched for dilution relative to threshold concentration), plotted for glomerular tufts and respective daughter TCs and MCs (n = 5 OBs, 4 mice) showing no systematic change in excitation ratio with concentration.

Glomerular MT GCaMP signals showed concentration- and odorant-dependent dynamics similar to those of OSNs or glutamate signals, with longer-latency response onsets (Δacid = 86 ± 12 ms, *p* = 6E-6, n = 16 pairs, Mixed-effects pairwise comparisons) and delayed ITA waveforms for the ester derivative of each carboxylic acid (Fig. 7B-D). MT dendrites also showed delayed responses to aldehyde and alcohol derivatives (Fig. S7A) (Aldehydes: Δacid = 254 ± 36 ms, *p* = 5E-5, n = 8; Alcohols: Δacid = 120, 127 ms, n = 2).

Glomerular MT signals reflect the summed postsynaptic dynamics across tufted cells (TCs) and mitral cells (MCs) and reflect a mixture of synaptically-driven and backpropagating action potential Ca^2+^ influx (Charpak et al., 2001). To assay spike output from individual MT neurons, we imaged from the somata of TCs or MCs innervating the glomerulus of interest; we refer to the cohort of cells innervating a glomerulus as ‘daughter’ cells (Dhawale et al., 2010; Karadas et al., 2026). Daughter TC somata were imaged <50 µm below the primary glomerulus and were visually confirmed by tracing the primary dendrite from the soma to the glomerulus using odorant-evoked ΔF signals; in fortuitous preparations we could image GCaMP signals in the ‘parent’ glomerular tuft and the somata of daughter TCs simultaneously in a single imaging plane (Fig. 7E). Up to 9 daughter TC somata innervating the parent glomerulus were imaged in the same field of view; as reported previously (Karadas et al., 2026), daughter TCs showed highly-correlated responses to odorant stimulation, and so were averaged together to improve the signal-to-noise ratio for latency measurements. GCaMP signals showed similar-latency response onsets as glomerular signals but were less clearly modulated by individual inhalations (Fig. 7F, G), possibly due to slower Ca^2+^ dynamics in the soma, although such modulation was apparent after inhalation-triggered averaging (Fig. 7H). As with the glomerular imaging, TC somatic signals showed delayed excitatory responses to the ester derivative of each primary acid (Fig. 7G, I) (*p* = 1E-4, Mixed-effects pairwise comparisons). Delayed onsets for successive inhalations of odorant were also apparent in the ITA waveform (Fig. 7H).

To identify daughter MCs, we first confirmed that the primary odorant at its near-threshold concentration evoked Ca^2+^ signals in a single glomerulus in the field of view (e.g., Fig. 7A), then moved the imaging plane to the mitral cell layer, ∼ 150 - 400 µm below the glomerulus (Fig. 7J). MCs showing significant excitatory responses to the carboxylic acid at the diagnostic (perithreshold) odorant concentration were designated daughter MCs (see Methods). We then compared responses for acid-ester pairs presented at 10 - 100x the perithreshold concentration. The number of daughter MCs simultaneously imaged in a field of view ranged from 3 to 9.

MC somatic GCaMP8f signals also showed delayed excitatory responses to the ester derivative of each acid tested (Δacid, onset latency: 146 ± 57 ms, *p* = 0.044, Mixed-effects pairwise comparisons, n = 7 parent glomeruli), and showed delayed ITA waveforms for successive inhalations (Fig. 7K, L, M). Ester-acid latency differences in glomerular signals, TCs and MCs decreased with increasing odorant concentration but remained longer for the ester at all concentrations (Fig S7C). Overall, these results indicate that timing relationships between carboxylic acids and their ester derivatives observed at the level of OSN input appear essentially unchanged at the level of OB output from tufted or mitral cells.

As implied by the analyses above, delayed ester-evoked inputs to acid-sensitive glomeruli did not emerge as suppressive responses at the level of TC or MC somata. Suppressive responses were evident in neighboring glomeruli and in adjacent (i.e., non-daughter) MCs in the same field of view (Fig. 7K, Fig. S8D, E), indicating that inhibitory circuits were engaged by odorant stimulation. However, even in cases where suprathreshold ester stimulation evoked shorter-latency excitation in neighboring glomeruli, responses in the primary glomerulus and its daughter cells consisted of delayed excitation rather than suppression (Fig. S8F).

To screen for inhibition of delayed ester-evoked inputs more systematically, we considered that timing-dependent inhibition by OB circuitry could manifest as a reduction in excitation at the level of MT cell output rather than outright suppression of baseline firing. To test for either possibility, we measured the ratio of excitation (non-negative ΔF/F) over the first second of odorant presentation for acid-ester pairs presented at threshold-matched concentrations (10x threshold and 10 - 100x perithreshold), measured from the parent glomerulus or the same daughter TCs and MCs, respectively, in a given experiment (Fig. 7N). Ratios were expressed as ΔF/F_ester_ : ΔF/F_acid_. Using this metric, ratios less than 1 reflect smaller mean activity in response to the ester, with values greater than 1 reflecting larger mean activity (ratios were computed over the first second of stimulation). The emergence of timing-dependent inhibition at the level of mitral or tufted cell output should appear as a decrease in ester : acid ratio in TC or MC somatic signals, relative to that of the parent glomerulus (Fig. 7N). Instead, there was no significant change in ester : acid excitation ratio for MC somata (*p* = 0.49, n = 21, Wilcoxon Signed Ranks), while TC somata showed an *increase* in excitation relative to their parent glomeruli (*p* = 0.004, n = 22; Wilcoxon Signed Ranks) (Fig. 7O). We also asked if there was a change in ester : acid excitation ratio as a function of concentration, as suprathreshold concentrations recruit increasing interglomerular inhibition (Banerjee et al., 2015; Economo et al., 2016; Karadas et al., 2026). In replicates of the tiglic acid - methyl tiglate identified glomerulus (n = 5), we observed no consistent change in ester : acid excitation ratio in glomerular tufts, TC or MC somata (Fig. 7P). Overall, these results do not support the prediction that inhibitory circuitry in the OB suppresses delayed inputs - at least those driven by odorant metabolism. Instead, they suggest that the altered chemical tuning and response dynamics resulting from nasal odor metabolism are relayed to higher-order olfactory areas.

## Discussion

Metabolism of odorants by xenobiotic enzymes in the nasal epithelium is a well-established phenomenon whose primary function is to facilitate the removal of odorant molecules from mucus and epithelial tissue (Heydel et al., 2013). This ‘housekeeping’ role has not typically been thought to substantially impact the initial encoding of odor identity, despite a few select reports in mice (Nagashima and Touhara, 2010; Kida et al., 2018). The collective results from the present study suggest that odorant metabolism broadly and profoundly shapes odor representations, and does so over a time-scale relevant for odor perception. Incorporating rapid odorant metabolism into a model of how inhalation drives OSN activation explains the diversity of OSN response dynamics more completely than a simple sensitivity-based timing model, and explains what would otherwise appear to be complex tuning of OSNs to chemical features to which their cognate odorant receptors are insensitive. While odorant metabolism introduces the potential for confusion in the encoding of inhaled versus metabolism-generated odorants, we find that the inhalation-linked timing features of OSN responses can robustly discriminate these odorant sources, and that timing differences are relayed out of the OB to higher centers. These results suggest a novel role for inhalation-linked timing in odor coding: to disambiguate inhaled odorants from those generated internally. They also raise the possibility that delayed sensory inputs from metabolism- or other internally-generated odorants are not simply suppressed but instead processed in parallel to those from external odorants, such that they may mediate distinct adaptive responses to olfactory stimuli arising from different sources (Fig. 8).

**Figure 8.**
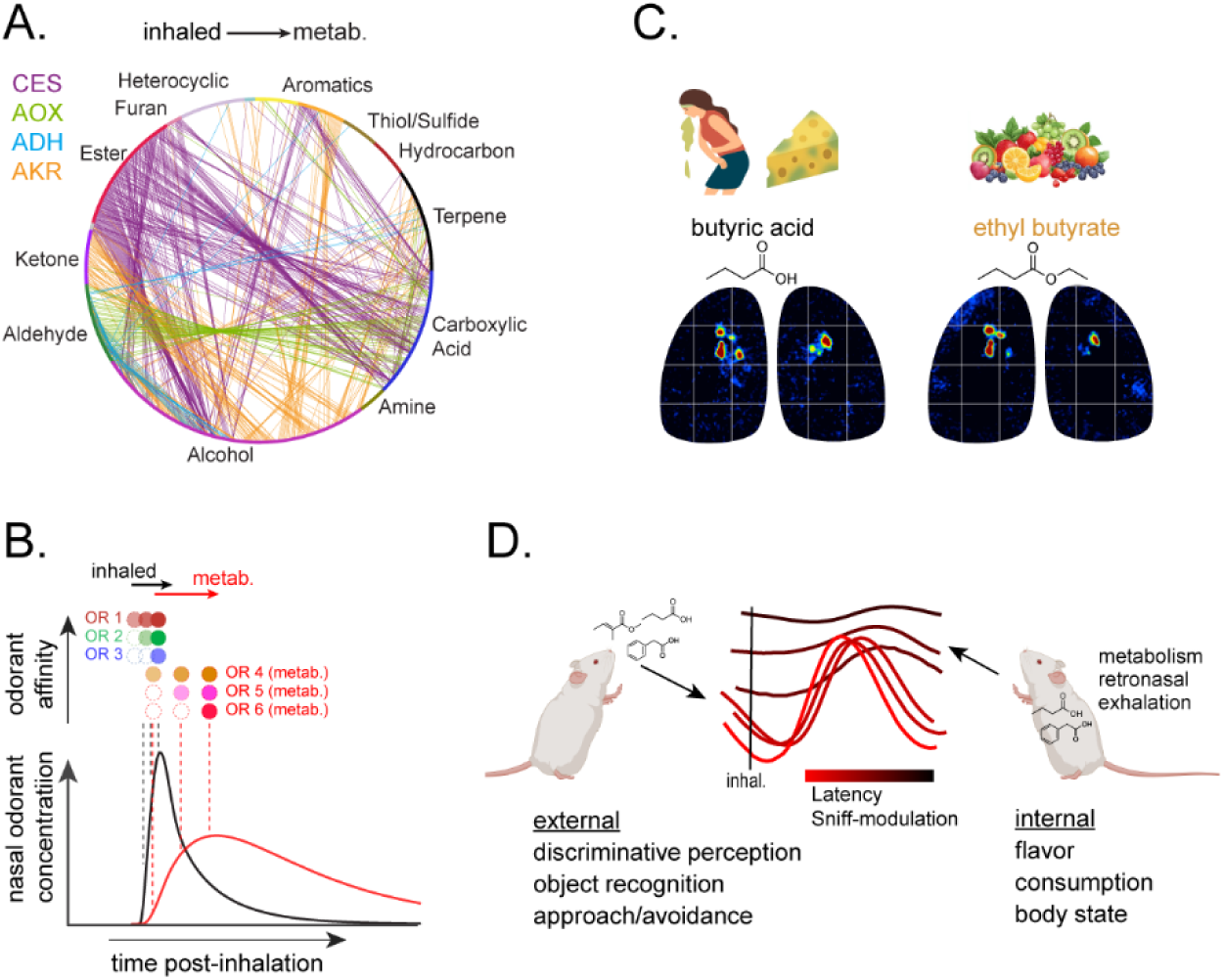
Revised model of inhalation-linked timing in odor representations and its potential role in disambiguating inhaled from internally-generated odorants. **A.** Network of predicted confusion between odorants and their metabolites, computed for four nasal metabolic enzymes on 632 odorants used in a survey of prior experimental studies (Wachowiak et al., 2025). CES, carboxylesterase; AOX, aldehyde oxidase; ADH, alcohol dehydrogenase; AKR, aldo-keto reductase. See Text for details. **B.** Revised sensitivity-timing model (compare to Fig. 5A), showing compressed latency range for odorant receptors (OR) activated by inhaled odorant, and delayed responses to odorant metabolites. Note that the most-sensitive metabolite ORs may be activated earlier than less-sensitive inhaled-odorant ORs. **C.** Example of near-identical glomerular representations for the ester, ethyl butyrate, and its acid metabolite, butyric acid, along with icons representing common odor descriptors for each compound. **D.** Potential scheme for disambiguating inhaled from internally-generated odorants on the basis of inhalation-linked timing.

We focused here on the actions of only a few classes of xenobiotic enzyme – those that convert esters and aldehydes into carboxylic acids - in explaining response features of OSNs that are not predicted by simple receptor-determined models. The impact of this metabolic pathway on odor representations was particularly noticeable due to the fact that a major subfamily of odorant receptor appears specialized to detect carboxylic acids (Cichy et al., 2019; Billesbølle et al., 2023) and the fact that their OSNs project to glomeruli in a distinct spatial domain of the OB (Bozza et al., 2009). By itself, cross-talk between representations of esters, aldehydes and carboxylic acids represents a major potential confound to olfactory function, as each class of odorant is abundantly represented in natural sources. However, these results likely represent only a sampling of the impact on odor representations mediated by the full repertoire of enzymes expressed in the nasal epithelium, which is large and diverse (Ibarra-Soria et al., 2014; Kornbausch et al., 2022). A predicted confusion matrix between odorants and their metabolites for just four enzymes with verified expression in nasal or oral mucosa (Bogdanffy and Taylor, 1993; Ijichi et al., 2019; Schwartz et al., 2021; Takaoka et al., 2022), using ∼600 odorants compiled from prior experimental studies (Wachowiak et al., 2025) (Fig. 8A), shows that odorant metabolism has the potential to introduce cross-talk between inhaled and enzyme-generated odorants across a large portion of olfactory chemical space, as has been proposed by others (Heydel et al., 2013; Kornbausch et al., 2022). Indeed, we observed delayed- and non-respiration-modulated responses to other odorant classes that may be targeted by other enzymes (e.g., 2’-OH acetophenone, Fig. S5) (Kornbausch et al., 2022). Further work is necessary to confirm that such responses are signatures of other metabolite-generated odorants, but if so, slow response dynamics could be used as a means to more fully characterize the extent of nasal metabolism in shaping early odor representations.

### Implications for timing-based strategies of odor coding

In the course of exploring how odorant metabolism impacts OSN response dynamics we characterized the relationship between odorant concentration, relative sensitivity and response timing in several functionally-identified glomeruli, using odorants to which they are exceptionally sensitive, which we nominally call their ‘primary’ odorants. These experiments provided a systematic test of predictions of the sensitivity-latency model of OSN response dynamics using multiple glomeruli matched to their primary odorants and tested within a naturalistic range of odorant concentrations (Wachowiak et al., 2025). We confirmed two foundational predictions of this model: 1) response latencies of glomeruli decrease with increasing odorant concentrations, and 2) relative latencies across glomeruli reflect their relative sensitivities to an odorant (Wilson et al., 2017; Giaffar et al., 2024). However, these predictions held true under a limited range of conditions. Concentration-dependent changes in response latency for primary odorants (carboxylic acids or ketones) had a dynamic range of ∼ 50 ms over up to three log units of concentration change (as measured with the fast reporter iGLuSnFR3; Fig. 4F, S4), and sensitivity-based timing differences (i.e., latency differences between the most-sensitive and less-sensitive glomeruli) ranged from a few tens of milliseconds to nonexistent, depending on glomerulus identity and odorant concentration. Even these modest sensitivity-latency relationships disappeared during repeated inhalations of odorant. The relatively small dynamic range of timing differences and the lack of robustness across different modes of odorant sampling constrain the use of timing-based coding strategies to achieve a concentration-invariant representation of odor identity or to mediate odor object recognition, as has been proposed (Hopfield, 1995; Wilson et al., 2017; Bolding and Franks, 2018; Gill et al., 2026; Karadas et al., 2026).

By contrast, odorants subject to nasal metabolism (e.g., esters, aldehydes) evoked responses with latency differences of 50 - 200 ms relative to other glomeruli (or other odorant-evoked responses), showed a larger dynamic range across concentrations, and maintained these timing differences across repeated inhalations of odorant. These results suggest that perireceptor processes such as nasal metabolism are a larger - and more robust - driver of temporal response patterns than strictly receptor-determined mechanisms (Fig. 8B). Notably, the range of timing differences attributable to odorant metabolism aligns with the range of temporal patterns reported in previous studies of odorant-evoked activity in the mammalian olfactory system (Chaput, 1986; Spors et al., 2006; Wachowiak, 2010; Short and Wachowiak, 2019; Moran et al., 2021b). Esters and aldehydes are commonly-used odorants in mammalian and nonmammalian model systems, constituting ∼25% of the ∼600 compounds compiled in a recent meta-analysis (Wachowiak et al., 2025). Thus, much of the temporal diversity that has been the focus of previous studies may reflect odorant metabolism.

We also found that esters activated class I OR-expressing, acid-sensitive OSNs at lower concentrations - but longer latency - than ester-selective glomeruli, leading to violations of the expected relationship between sensitivity and latency predicted by a receptor-determined model (Hopfield, 1995; Wilson et al., 2017). This apparent anomaly could be explained by odorant metabolism if class I ORs are more sensitive to their carboxylic acid ligand than are class II ORs to the parent ester. This sensitivity difference need not be large, as our simulations suggested that acid concentrations produced by ester metabolism can reach ∼20% of the peak inhaled ester concentration. The higher water-solubility of carboxylic acids relative to their parent ester would also favor preferential concentration of the acid product in the mucus. It will be interesting to explore whether sensitivity-latency violations are apparent with aldehydes, additional esters or odorant pairs linked by other enzymatic pathways. In addition, our findings with respect to the esters characterized here present an opportunity to further test models of odor identity perception using stimuli whose identity- and timing-based representations change predictably with concentration.

### Disambiguating inhaled from metabolism-generated odorants

The diversity and abundance of enzyme expression in the nasal epithelium suggests that the adaptive value of facilitating removal of inhaled odorants from the upper airway outweighs the potential confounds to odor identity coding. Nonetheless it seems crucial that the nervous system disambiguate sensory signals reflecting inhaled odorants from those generated by their metabolites. For example, in the natural environment carboxylic acids are common products of bacterial action and organic decomposition (i.e., spoilage or fermentation) (St. Angelo et al., 1996; Ng et al., 2023) and at least some acids elicit innate avoidance (Kobayakawa et al., 2007; Cichy et al., 2019), while esters are commonly produced by cellular metabolism in flowers and fruits (Beekwilder et al., 2004; Ceccoli et al., 2024), often signaling a nutritive food source (Fig. 8C).

Inhalation-linked timing provides a robust neural feature by which to disambiguate these signals: initial onset latencies and the degree of modulation over successive inhalations supported discrimination of responses to a carboxylic acid versus its metabolic parent ester with high accuracy. Timing also effectively separated glomerular responses evoked by direct inhalation of an ester versus its presumed acid metabolite, activated in the same trials. Multiple lines of evidence suggest that the earliest-activated inputs to the olfactory system preferentially drive odor perception (Rinberg et al., 2006; Wesson et al., 2008; Chong et al., 2020); this phenomenon - termed ‘primacy’ (Wilson et al., 2017; Giaffar et al., 2024) - has been proposed to mediate concentration-invariant odor recognition (Hopfield, 1995) and odor learning (Gill et al., 2026). We suggest that perceptual ‘primacy’ is also adaptive for avoiding confusion between inhaled odorants and their nasal metabolites, and the present results motivate further experiments testing this hypothesis with behavioral and neural measures.

We initially hypothesized that the separation between inhaled and metabolism-generated odorants is achieved by circuitry within the OB, as a recent study reported that OB circuitry suppresses MT cell responses to delayed sensory inputs over the same time-scale (Karadas et al., 2026). However, while we observed abundant evidence of odorant-evoked suppression in MT cell signals, timing differences between carboxylic acids and their ester derivatives - present at the level of OSN input and in MT cell glomerular tufts - persisted in MT cell excitatory responses. We found no evidence for an emergent suppression or even a selective loss of excitatory drive for (slow) ester versus (fast) acid-evoked inputs. One explanation for the discrepancy between these two studies may lie in the magnitude and extent of inhibition elicited. In the previous study (Karadas et al., 2026), inhibition was evoked with approximate odorant concentrations 3 - 5 log units higher than those used here (2E-3 - 2E-2 total dilution in (Karadas et al., 2026) versus ∼2E-7 - 2E-5 in the present study), driving OSN input to more glomeruli and more extensively activating the lateral inhibitory network within the OB (Banerjee et al., 2015; Economo et al., 2016). Such concentrations rarely, if ever, occur in natural settings (Wachowiak et al., 2025). Delayed responses in Karadas et al. also were consistently smaller in magnitude than early responses, as measured in MT cell dendrites (Karadas et al., 2026). Here, we compared odorant-evoked responses of similar magnitude (as measured directly from OSN terminals) but varying in onset latency, allowing for dissociation of timing- and intensity-dependent effects. Another possibility is that timing-dependent inhibition may selectively occur between particular glomeruli, as reported previously (Economo et al., 2016), obscuring our ability to detect it. While further exploration of the determinants of MT cell suppression is necessary, the key takeaway from our results is that metabolite-driven sensory signals are not filtered by OB circuits. Instead, timing-based features distinguishing inhaled from metabolism-generated odorants persist at the level of MT cells and are relayed to olfactory cortical areas.

Piriform cortex is a likely candidate for separating signals arising from inhaled versus metabolism-generated odorants. Piriform circuits mediate suppression of MT inputs that are delayed on the 50 - 100 ms time-scale typical of metabolite-driven MT activation and may also suppress responses to tonic MT activity (Stokes and Isaacson, 2010; Bolding and Franks, 2018). The timing-based suppression of MT signals by piriform circuitry has been proposed to mediate concentration-invariant odor identity coding (Bolding and Franks, 2017; Stern et al., 2018). Our results suggest an alternative function for this process, which is to disambiguate neural signals arising from inhaled odorants and their metabolites (Fig. 8D). A prediction arising from this hypothesis is that principal neurons in piriform cortex will not show co-tuning to carboxylic acids and their ester derivatives, while such co-tuning is uniformly seen in MT cells. A previous study found systematic differences between MT cells and piriform neurons in the representation of physicochemical feature space (Pashkovski et al., 2020), and we speculate that odorant-metabolite relationships will provide additional explanatory power in further understanding this transformation.

### Implications for odor perception

Given the evidence that sensory signals from odorant metabolites are relayed to olfactory cortical areas, it is intriguing to consider the extent to which such signals shape the structure of odor perception. A number of studies have reported instances in which odorant metabolism likely shapes odor perceptual qualities in humans (Kornbausch et al., 2022; Shirai et al., 2023; Ye et al., 2024). Previous analyses have reported that intracellular metabolic relationships may partially explain perceptual similarities between odorant compounds (Chee-Ruiter, 2000; Qian et al., 2023); these studies considered metabolic processes within an odor *source* as driving co-occurrence of different compounds. Nasal odorant metabolism, by contrast, generates co-occurrence of odorants within the *receiver* and does so independent of odor source, suggesting that odorants linked by nasal metabolism may be especially likely to share perceptual descriptors if co-occurrence is an important factor driving odor quality perception.

A final consideration is the possibility that sensory signals arising from odorant metabolism are not simply filtered on the basis of their distinct timing but instead are represented and processed in parallel to signals from inhaled odorants (Fig. 8D). Such a strategy may generalize beyond nasal metabolism to include the differential processing of external versus internally-generated odorants, which can carry crucial information about body state and recent food consumption (Blankenship et al., 2019; Mori and Sakano, 2022), potentially mediating distinct but adaptive behavioral responses to such signals.

### Limitations of the study

One important limitation of this study is that we have not directly demonstrated that the co-tuning relationships and delayed temporal responses in acid-sensitive glomeruli are due to nasal odorant metabolism. While rapid nasal metabolism is the most parsimonious explanation for our results, and the phenomenon is well-established in both rodents and humans (Nagashima and Touhara, 2010; Heydel et al., 2019; Robert-Hazotte et al., 2019; Robert-Hazotte et al., 2022), this conclusion would be strengthened by pharmacological blockade or genetic knockout of the enzymes involved, in conjunction with functional measures. Previous attempts at pharmacological manipulation of odorant - evoked responses in vivo have yielded complex and inconsistent effects (Nagashima and Touhara, 2010), and the relevant enzymes (e.g., carboxylesterase, aldehyde dehydrogenase) are only partially effective at reducing nasal enzymatic activity (Stanek and Morris, 1999; Nagashima and Touhara, 2010). Genetic manipulations are a promising strategy, but are difficult given that most conversion pathways are mediated by multiple enzyme subtypes, each encoded by a distinct gene, with potentially overlapping substrate specificities (Satoh and Hosokawa, 2006; Kornbausch et al., 2022). Nonetheless, genetic removal of key enzymes would allow for a broader exploration of the impacts of odorant metabolism on odor coding, relative to the contribution of receptor-determined mechanisms.

Our study was limited to imaging activity in, or from, glomeruli of the dorsal OB, and especially from acid-sensitive glomeruli in the class I-recipient domain (Bozza et al., 2009); thus, sensitivity-timing relationships characterized here may be missing non-dorsal glomeruli with higher sensitivity to ester or aldehyde ligands. This caveat applies to many other recent studies characterizing odor coding strategies, especially those relying on optical approaches which have, with few exceptions, focused on the same area of the dorsal OB (Rubin and Katz, 1999; Spors et al., 2006; Ma et al., 2012; Chae et al., 2019; Burton et al., 2022; Karadas et al., 2026). Our imaging approach also limits interpretation of activity patterns in postsynaptic neurons, as even the fastest-available GCaMP8f reporter is limited in its ability to report fine-scale timing and changes in spiking activity over the full dynamic range of mitral and tufted cells. Electrophysiological recordings in awake mice would thus be useful in future studies.

## Materials and Methods

### Animals

Mice were used from the following genetically-modified strains, obtained from the Jackson Laboratories: OMP-Cre, JAX Stock #006668; TIGRE2-jGCaMP8m-IRES-tTA2-WPRE (’TIGRE2-GCaMP8m’), Jax# 037718; Tbet-Cre, JAX #024507. The data presented and re-analyzed from (Burton et al., 2022), and two of 6 mice used for the data presented in Fig. 2B, were collected in heterozygous crosses of OMP-IRES-tTA (Jackson Laboratory stock #017754) (Yu et al., 2004) and tetO-GCaMP6s (Jackson Laboratory stock #024742) (Wekselblatt et al., 2016) mice aged 2-6 months (OMP-IRES-tTA x tetO-GCaMP6s mice). Or52h2-IRES-mKate2 mice were generated at the University of Utah Gene Targeting Core by inserting an *IRES*-*mKate2* cassette downstream of the CDS of the OR gene using *Easi*-CRISPR. Or52h2-IRES-mKate2 mice are available from the investigators on request. Mouse lines were maintained in a breeding colony at the University of Utah and were crossed as described in the Text. Experiments used male and female mice from 3 - 6 months in age. Mice were housed up to 4/cage and kept on a 12/12 h light/dark cycle with food and water available ad libitum. All procedures were carried out following the National Institutes of Health Guide for the Care and Use of Laboratory Animals and were approved by the University of Utah Institutional Animal Care and Use Committee.

### Viral Vectors

Expression of iGluSnFR variants and GCaMP8f in M/T cells was achieved with Cre-dependent expression of viral vectors in the OB of Tbet-Cre mice. Viral vectors used were: AAV5-Flex-GCaMP8f (pGP-AAV-syn-FLEX-jGCaMP8f-WPRE, Addgene #162379); AAV1-FLEX-iGluSnFR3 (pAAV.hSyn.FLEX.iGluSnFR3.v857.GPI.codonopt, Addgene #17581); AAV5-FLEX-SF-iGluSnFRA184S (pAAV.hSyn.FLEX.SF-iGluSnFR.A184S, ViGene). Virus injection (300 - 500 nL, ∼250 μm depth, 1:2 - 1:10 dilution) was done using pressure injections and beveled glass pipettes, as described previously (Short and Wachowiak, 2019), during the chronic window surgery.

### Awake, head-fixed imaging

For two-photon imaging in awake, head-fixed mice we prepared a headbar and chronic imaging window implanted over both OBs under isoflurane anesthesia, as described in (Karadas et al., 2026). The window consisted of a 3 mm diameter coverslip that spanned the midline and included a field of view of approximately the anterior and medial half of each OB. Animals were allowed two weeks to recover from surgery and allow for window clearing and virus expression. Animals were then acclimated to head-fixation on a circular treadmill (Nakayama et al., 2022) over the course of 4 - 6 days, and subject to at least one 30-minute session including odorant presentation prior to data collection. Imaging sessions lasted from 30-60 minutes per day, over a span of 5 - 30 days.

For widefield imaging in awake, head-fixed mice, a headbar was implanted and the skull overlying the OB was exposed, cleaned with hydrogen peroxide (1%) and 70% ETOH and sealed with Kwik-Sil (WPI). After recovery from surgery mice were acclimated to head-fixation and odorant presentation. After acclimation, mice were anesthetized with isoflurane, the Kwik-Sil removed and the bone overlying nearly the full extent of each dorsal OB thinned to transparency using a dental drill (Osada USA) at low speed. A thin layer of cyanoacrylate was applied to maintain transparency and both OBs were covered with a single 4 mm coverslip sealed in place with optical adhesive (NOA 61, Norland Products, Cranbury, NJ). Data collection in the awake mouse began the following day, and continued for 3 - 5 days or until the underlying bone regained opacity.

### Data collection

In most experiments, two-photon imaging was carried out with a Neurolabware in vivo microscope two-photon microscope (Sutter Instruments or Neurolabware) coupled to a Chameleon Ultra pulsed Ti:Sapphire laser (Coherent) at 940 nm, controlled by Scanbox (Neurolabware) software. Imaging frames were acquired with bidirectional resonance scanning at 30 Hz. For two-photon imaging of GCaMP8m in Or52h2-IRES-mKate2 mice, a dual-color two-photon system was used, consisting of a Sutter Instruments scanning microscope, a Mai Tai HP Ti:Sapphire laser (Spectra-Physics) at 920 nm for GCaMP imaging and a Fidelity-2 fixed-wavelength (1060 nm) laser for mKate2 visualization, with images collected using unidirectional resonance scanning at 15.2 or 15.5 Hz under the control of ScanImage (Vidrio).

For widefield imaging, epifluorescence was collected through a 4x, 0.28 N.A. air objective (Olympus) at 256×256-pixel resolution and 50-Hz frame rate using a back-illuminated CCD camera (NeuroCCD-SM256; RedShirt Imaging) under the control of SM-Turbo software. To minimize hemodynamic artifacts that were prominent in the awake mouse, a dual-wavelength LED illumination system was designed to allow collection of fluorescence emission at the isosbestic excitation point for hemoglobin on alternate frames. Custom Arduino code (kind gift of D. Storace, Florida State Univ) used the camera frame clock to trigger a 470 nm LED or 405 nm LED (M405L4, M470L3, Thorlabs) on alternate frames. LED light was collimated using an aspheric condenser lens (ACL2520U, Thorlabs) and merged into the same illumination path within a 1”-diameter cube (DFM1, Thorlabs) containing a dichroic mirror (DMSP425R, Thorlabs) then passed through a multi-band bandpass filter (69000x, Chroma Technology) before transmission to the objective. Fluorescence emission was collected through the standard Olympus fluorescence turret using a GFP4050a filter cube (Semrock) with emission filter removed. The intensity of each LED was adjusted manually to achieve roughly equal fluorescence emission before data collection. Strobed frames were collected and digitized in sequence. Subsequently, 470 nm : 405 nm ratio images were computed from successive frame pairs on a pixel by pixel basis. Odorant-evoked fluorescence changes are thus expressed as change in ratio (ΔR).

Odorants were presented as square pulses lasting 4 - 8 seconds and were typically repeated at the same concentration over 6 - 8 trials, with imaging data collected continuously throughout the series of repeats. An interstimulus interval between 15 - 30 sec was chosen to avoid adaptation between presentations, depending on odorant identity and concentration. Respiratory signals were acquired via tubing connected to a flow sensor (AWM3150V, Honeywell) positioned at the external nares on the side opposite that being imaged, digitized at 1 kHz and synchronized with the imaging frame clock.

### Odorant preparation and presentation

Odorants used are listed in Table S1. In most experiments odorants were presented using a custom olfactometer (a.k.a, ‘Turbulator’) that allowed for precise temporal control of odorant onset and offset. The device used a novel design that achieved rapid and high dilution of odorant vapor in free space and generated reliable odorant pulses with a 90% rise-time of ∼50 ms, with a minimal (<0.2%) change in net air flow to the mouse (Supplemental Figure S3). The design also enabled linear control of dilution levels over a four-fold range and choice of eight odorant channels with no common flow path, allowing for rapid switching between odorant channels and minimal chance for cross-contamination between channels. Fast odorant mixing and dilution occurred via an inexpensive, commercially-available Venturi-based fluid mixing device (BEX mini tank mixing eductor, 0.059” orifice, item# 62386, Unites States Plastic Corp.). Saturated vapor from glass odorant reservoirs with teflon inflow and outflow ports (Diba Omnifit Q Series GL45 cap, Cole-Parmer) was directed to dedicated T-connectors (Clippard 1/16” swivel tee, ST0-2002) arrayed around the eductor inlets and released into the eductor by closing a normally-open valve attached to vacuum. The small dead volume of the T-connector allowed for rapid odorant onset and offset that was shorter than the typical inter-inhalation interval in an awake, resting mouse (Fig. S4). Speed and precision of timing was characterized with a photoionization detector (mini-PID, Aurora Scientific). Amount of dilution achieved by the eductor relative to saturated vapor was 0.2%, measured at the position of the mouse nose (5 cm from the eductor nozzle) under standard operating parameters (see Fig S3 legend). Vapor-phase dilution ranged from two-fold less to two-fold higher depending on flow rate through the odorant reservoir (Fig. S3B). For the characterization of glomerular specificity across ester and aldehyde derivatives (Fig. 2B-E), odorants were presented using a high-throughput odor delivery device (aka, ‘Odor Gun’), described previously (Burton et al., 2019), which used the same eductor-based air dilution principle and achieved a vapor-phase dilution of ∼1.3%. Additional dilutions (by factor of ten, typically, 1E-2 to 1E-6) were achieved by diluting pure odorant in an odorless solvent (caprylic/capric medium-chain triglyceride oil (MCT), C3465, Spectrum Chemical Mfg. Corp.). Odorant concentrations are reported in the Text and Figures as total dilution, accounting for liquid dilution and subsequent air-phase dilution. This reporting is based on ideal (linear) assumptions of vapor pressure after dilution in liquid (i.e., Raoult’s Law) and the subsequent air-phase dilution by the eductor. While many odorants deviate from ideal behavior as described by their Henry’s Law coefficient (Cometto-Muñiz et al., 2003; Jennings et al., 2023) (which is uncharacterized for the MCT oil solvent), this deviation predominates at more-concentrated liquid dilutions, with subsequent dilutions in solvent increasingly approaching linearity. Thus, relative changes in odorant concentration can be assumed to accurately track liquid dilutions at the high degrees of dilution used in most experiments. Estimated delivered molar concentrations, calculated from the odorant’s vapor pressure and ideal assumptions, are given for reference dilutions in Table S1 and are included to highlight differences in concentration among different odorants and facilitate comparison with other studies.

### In vitro OR - ligand screening

GloSensor cAMP assays (Promega) were used to measure intracellular cAMP levels in Hana3A cells downstream of odorant receptor activation, as previously described (Mainland et al., 2014; Zhang et al., 2017). Hana3A cells were cultured in minimum essential medium (MEM; Corning) supplemented with 10% fetal bovine serum (FBS; Gibco), penicillin-streptomycin, and amphotericin B (Ikegami et al., 2020). Cells were seeded the day before transfection at a density of 8 × 10⁶–1 × 10⁷ cells/mL. For each 96-well plate, 1,000 ng pGloSensor-20F plasmid (Promega), 500 ng RTP1S plasmid, and 7,500 ng rho-tagged odorant receptor in the pCI mammalian expression vector (Promega) were co-transfected using 20 µL Lipofectamine 2000 (Invitrogen) in MEM supplemented with 10% FBS. After 18–24 h, transfection media were replaced with 25 µL of 2.6% GloSensor substrate (Promega) diluted in HEPES-buffered solution supplemented with D-glucose, and plates were incubated at room temperature in the dark for 2 h prior to measurement.

For in vitro odorant stimulation, 1 M stock solutions were prepared in ethanol and diluted on the day of the assay into Hank’s Balanced Salt Solution (HBSS) supplemented with D-glucose to achieve the indicated final concentrations. A separate saturated stock solution was prepared as the highest starting concentration. Half-log serial dilutions were then generated in HBSS with D-glucose to produce seven odor concentrations and a no-odor control, as previously described (Saito et al., 2009; Mainland et al., 2014; Zhang et al., 2017).

Luminescence was measured for 15 min immediately following odorant stimulation using a CLARIOstar^Plus^ plate reader (BMG LABTECH). cAMP accumulation was quantified by integrating luminescence values across measurement cycles (expressed as area under the curve, ‘AUC’), as previously described (Zhang et al., 2017). Responses were normalized to empty vector controls and baseline luminescence (measured prior to odor addition). Dose–response curves were fitted using a three-parameter logistic regression model in Prism 10.6.1 (GraphPad Software).

### Analysis of chemical feature space

To assess odorant relationships based on physicochemical features and xenobiotic metabolism pathways, we used the Python package RDKit (https://www.rdkit.org/). To visualize and quantify odorant similarities (e.g., Fig. 2C, D), an odorant physicochemical feature space was constructed from 4989 odorous compounds in the combined GoodScents/Leffingwell dataset (Lee et al., 2023), as provided by the OpenPOM github site (https://github.com/ARY2260/openpom). This list was supplemented with 12 additional compounds from our experimental odorant panel. RDKit was used to generate molecular fingerprints for each compound from its SMILES string; fingerprints were encoded as MACCS keys, a binary representation indicating the presence or absence of 166 chemical features (Durant et al., 2002). Principal component analysis (PCA) embeddings were computed from MACCS fingerprints of the chemical space dataset. For display (Fig. 2C), a three-dimensional scatter projection of the first three PCs was visualized using OriginLab (Origin(Pro) 2026, OriginLab Corporation, Northampton, MA, USA); for visualization purposes, 3615 odorants were displayed instead of the full set.

To quantify physicochemical diversity among odorant groups (Fig. 2D), variance from the group centroid was defined as the variance in the distribution of Euclidean distances between individual odorants and the centroid of their respective group, calculated in the first four PCs, which included 39% of total physicochemical variance.

To visualize the potential for confusion between odorants and their metabolites (Fig. 6H), metabolic reaction products across odorant space were computed based on known reactions of four xenobiotic enzymes: carboxylesterase, aldehyde oxidase, alcohol dehydrogenase and aldo-keto reductase. These enzymes were selected from a larger set detected in rodent olfactory epithelium (Heydel et al., 2019; Boichot et al., 2023) due to their straightforward chemical reactions with particular chemical substrates. Substrate odorants and their predicted reaction products were identified using RDKit’s rdChemReaction module. For each enzyme, a general reaction pattern was defined using SMARTS notation (Daylight Chemical Information Systems Inc; https://www.daylight.com/dayhtml/doc/theory/theory.smarts.html), which specifies the structural changes a molecule can undergo. These reactions were then applied to a compilation of 632 odorants taken from previous experimental studies (Wachowiak et al., 2025). In total, 341 unique reactants were predicted to yield 354 unique products. When reactants and products were combined and duplicates removed, the final set, consisting of 660 unique odorants, was used to construct a circular network graph (907 nodes) using the python package ‘NetworkX’ (Hagberg et al., 2008), with lines connecting enzyme target and product odorant nodes (Fig. 8A).

### Real-time measurement of odorant metabolism by olfactory mucosa

The speed of ester metabolism by mouse olfactory mucosa was measured directly from excised tissue using proton transfer reaction - time of flight - mass spectrometry (PTR-TofF-MS). Esters used were methyl tiglate (CAS# 41725-90-0), methyl 2-methylbutyrate (CAS# 868-57-5), methyl propionate (CAS# 554-12-1) and isopropyl tiglate (CAS# 1733-25-1). We measured production of the alcohol hydrolysis product (methanol, CAS# 67-56-1; isopropanol, CAS# 67-63-0), as carboxylic acid detection was not sufficiently sensitive with the given PTR-MS configuration. All chemicals were purchased from Sigma Aldrich (Sigma Aldrich, St Quentin Fallavier, France).

Experiments were performed on 3 - 6 months male c57bl6 mice bred and housed in the animal facility of the Plateforme de Zootechnie at University of Dijon, France. Mice were housed in ventilated rack cages, with enrichment cell sizzle (SAFE, Augy 89) fed ad libitum with standard food from SAFE (A03-10 pour breeding, next A04-10). After decapitation, olfactory epithelium (OE) was carefully dissected to avoid contamination with respiratory epithelium and immediately frozen (cryotubes in liquid nitrogen) until ready for use. We used a total of 15 mice; 5 mice were used per molecule and per experiment.. All experimental protocols were conducted in accordance with ethical rules enforced by French law, and were approved by the local Ethical Committee of the University of Burgundy (Comité d’Ethique de l′Expérimentation Animale Grand Campus Dijon; C2EA grand campus Dijon N°105), and by the French Ministère de l′Education Nationale, de l′Enseignement Supérieur et de la Recherche under the no. 3504.

Measurements were conducted with a proton transfer reaction-mass spectrometer instrument equipped with a Time-of-flight analyzer (PTR-ToF 8000, Ionicon Analytik, Innsbruck, Austria). Data were recorded with the TOF-DAQ software in the range m/z 0 to 227.42 at one spectrum per 50 ms or one spectrum per 500 ms. The set up was identical to that previously validated and published (Robert-Hazotte et al., 2019). During the acquisition we focused on specific masses: m/z = 115.075 (protonated methyl tiglate), m/z = 116.079 (protonated isotope 13C methyl tiglate), m/z = 117.091 (protonated methyl-2-methyl butyrate), m/z = 118.085 (protonated isotope 13C methyl-2-methyl butyrate), m/z = 89.059 (protonated methyl propionate), m/z = 90.063 (protonated isotope 13C methyl propionate), m/z = 143.107 (protonated isopropyl tiglate), m/z = 144.110 (protonated isopropyl tiglate) and for the metabolites produced, m/z = 33.033 (protonated methanol) and m/z = 61.065 (protonated isopropanol). All measurements were carried out at a drift-tube pressure of 2.3 mbar, a drift-tube temperature of 80 °C and a drift-tube voltage of 390 V giving an electric field strength to number density ratio E/N of 116 Td (1 Td = 10^−^17 cm^2^.V) and H_3_O^+^ as reagent ion. The data were corrected for transmission and expressed in concentration (ppb) using primary ions H_3_O^+^ and (H_2_O)_2_H^+^ to account for primary ions fluctuations.

Odorant was drawn over excised olfactory mucosa using a flow control setup as described previously (Robert-Hazotte et al., 2019). Briefly, the flow configuration to the PTR-MS consisted of two independent, oven-thermostated circuits (control and experimental) (30 °C) controlled via a 6-way valve, allowing alternate delivery of odorants to either circuit (Supplemental Fig. S2A). Each circuit contained a glass trap: one received OE tissue from a single mouse and the other served as a control. Airflow was distributed via stainless-steel strainers and delivered to the PTR-MS through heated PEEK capillaries at 160 mL/min. Humidified zero-air from a Tedlar® gas bag maintained constant moisture, while a second bag contained aqueous odorant solutions (10 ppm in liquid). During experiments, background signals were first recorded in the control circuit, followed by timed introduction of odorants to both control and experimental circuits in a stepwise manner. Between runs, traps and tubing were cleaned to prevent odorant carryover. After the experimental run, the olfactory mucosa explant was heated at 80 °C for 15 minutes in a glass vial to inactivate enzymatic proteins, allowed to cool for 10 minutes, and then reintroduced into the system for analysis as a control for enzyme-dependent biotransformation.

The temporal delay (ΔT) between the PTR-MS signal of a parent odorant compound (A) and that of its enzymatically generated metabolite (B) was estimated using a relative rise-threshold approach combined with linear interpolation. This procedure was developed to provide sub-cycle temporal resolution beyond the nominal 50 ms PTR-MS acquisition interval while minimizing the influence of baseline fluctuations and transient signal noise, following principle commonly used in threshold-based signal onset detection (Kay, 1998). For each signal trace, the baseline was defined as the mean of the first ten data points within the analyzed response window, and the maximum as the highest intensity observed within the same window. A detection threshold was then calculated independently for each signal as the baseline plus 20% of the signal amplitude (maximum − baseline).

. The onset time of the parent compound (tA) was defined as the first crossing of the corresponding threshold during the rising phase of the signal. Because metabolite formation is expected to follow parent compound detection, the metabolite onset time (tB) was defined as the first threshold crossing occurring after tA. To reduce false-positive detections caused by noise or baseline fluctuations, threshold crossings were accepted only when they corresponded to a sustained rise of the signal rather than to an isolated fluctuation around the threshold. Threshold-crossing times were refined by linear interpolation between the last point below threshold and the first point above threshold. The minimum observable temporal delay was then calculated as Δt = tB − tA and reported in milliseconds. As Δt depends on a predefined relative threshold and on the temporal resolution of the acquisition system, it should be regarded as an operational estimate of the minimum observable delay between parent and metabolite appearance rather than as an absolute measure of the underlying enzymatic reaction kinetics.

Maximum methanol concentration produced by OE exposure to methyl tiglate was estimated by PTR-ToF-MS analysis from a calibration curve (y=169.44x, R^2^= 0.9814) obtained by the dilution of a 24.4 mg/l stock solution of methanol.

### Epithelial metabolism model

To simulate the dynamics of odorant and metabolite concentration changes during inhalation, an odorant transient derived from computational fluid dynamics (CFD) modeling of airflow in the rat nasal cavity was coupled to a model of transport and metabolism in the mucus and epithelium. The odorant transient was taken from a previous computational fluid dynamics (CFD) model of the rat nasal passage (Scott et al., 2014; Kim and Zhao, 2022; Wu et al., 2024), simulating concentration-change dynamics from a single inhalation at the at the anterior dorsal olfactory region. To model dynamics across successive inhalations the transient was repeated at 4 Hz (Fig. 3H). Odorant uptake at the air–mucus interface was handled through an effective boundary condition based on odorant solubility, using logP as a convenient estimator (Kurtz et al., 2004; Scott et al., 2014).

Following uptake, a one-dimensional, two-layer model was used to compute odorant transport and metabolism in the nasal lining. The domain includes a 30 μm mucus layer and a 70 μm epithelial layer. Diffusion of the parent compound (e.g., methyl tiglate) and its metabolite (e.g., tiglic acid) was modeled in each layer. Different diffusivities were assigned to the mucus and epithelial layers. Diffusivity in the mucus was estimated separately for the parent compound and its metabolite (*D*_1_ and *D*_2_, respectively) based on the Wilke–Chang method (Wilke and Chang, 1955), while diffusivities in the epithelium were reduced by a factor of 50 to reflect hindered transport in tissue.

Metabolism of the parent compound was described using Michaelis–Menten kinetics (Bogdanffy and Taylor, 1993; Corley et al., 2015). The parameters *V_max_* (maximum reaction rate) and *K_m_* (Michaelis constant) were taken from experimentally-determined values for carboxylesterase hydrolysis of vinyl acetate in the rat nasal mucosa (Bogdanffy and Taylor, 1993). The model accounts for both consumption of the parent compound and generation of its metabolite.

The governing equations are given in Eq. (1).

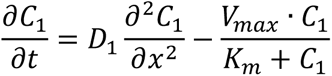

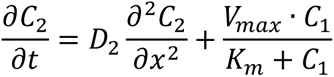

Where *C*_1_, *C*_2_ represent the concentration of the compound and its metabolite, respectively.

The model was implemented in MATLAB (MathWorks, Inc., Natick, MA) using the built-in solver *pdepe*. The spatial domain was discretized across both layers, and time-dependent simulations were driven by the odorant concentration profile obtained from the CFD model as the inlet boundary condition. This input was applied as a time-varying concentration using interpolation, while a zero-concentration condition was specified at the distal boundary. Layer-specific diffusivities and reaction terms were defined as piecewise functions of spatial location. The governing equations were solved simultaneously for the parent compound and its metabolite, accounting for diffusion and Michaelis–Menten kinetics.

The model used the following parameter values: Vmax, 3.7e-4 (mol/L/sec); Km, 3.33e-3 (mol/L); logP (methyl tiglate), 1.69; logP (tiglic acid), 1.4; D1, 7.8e-10 and D2, 8.6e-10 (m^2^/sec) in the mucus, with peak odorant concentration values in the air-phase as given in the Text.

### Imaging Data Analysis

#### Initial data processing pipeline

Data were initially processed using a custom GUI-based software written in Matlab (https://github.com/WachowiakLab/ImageAnalysisSoftware), as described previously (Moran et al., 2021b). For time-series analysis of two-photon data, regions-of-interest (ROIs) were chosen manually from the mean fluorescence image and further refined based on odorant response ΔF maps, then all pixels averaged within an ROI and time-series upsampled to 150 Hz (pchip function, Matlab) for further analysis. Respiration signals were low-pass filtered at ∼12 Hz and the times of each inhalation start and peak automatically identified using signal processing algorithms described previously (Moran et al., 2021b). For display in the figures, odorant response maps were generated using the change in fluorescence (ΔF) relative to the average of the 4 seconds preceding odorant onset values, after averaging all repeated odorant presentations. For display only, ΔF rather than ΔF/F values were used to minimize noise from nonfluorescent regions. Response maps were scaled as indicated in the figure and were kept to their original resolution (512 x 512 pixels) and smoothed using a Gaussian kernel with σ of 1 pixel.

#### Analysis of inhalation-linked activity dynamics

Inhalation-linked response timing features were measured from fluorescence traces of glomerular ROIs after aligning ROI traces to the time of inhalation, measured from the respiration signal. For initial onset latencies, traces from individual odorant presentations (3 - 6 successive trials) were aligned to the first inhalation after odorant onset, then averaged across trials. Because odorant presentation was not triggered by the respiration signal, trials in which odorant onset occurred within 50 ms of inhalation start were excluded from averaging, as were trials in which onset occurred during occasional bouts of high-frequency sniffing. After averaging, ROI traces were z-scored relative to the pre-odor baseline (1 - 2 seconds preceding odorant onset). Onset latency was defined as the time for the trace to reach a threshold level after which it must remain above threshold for 10 consecutive points (67 ms). A threshold of z = 3 was used for two-photon data; for widefield data the threshold was set to z = 4 - 6 due to the higher signal-to-noise ratio of responses. Signals failing to cross threshold within 1 sec of odorant onset were considered non-responsive.

For successive-inhalation analysis, inhalation-triggered average waveforms were constructed by averaging 400 ms-duration snippets of the ROI trace taken from the start of each inhalation occurring after the first 500 ms of odorant presentation, and analyzed as described in (Moran et al., 2021b). To avoid timing artifacts from preceding or later inhalations, only inhalations occurring at a frequency below 5 Hz (200 ms inter-inhalation interval) were included for averaging. Modulation Index (MI) was calculated from the ITA waveforms. For analysis of two-photon imaging data, MI was defined as the peak-to-trough amplitude of the ITA waveform divided by its mean ΔF relative to pre-odor baseline, then divided by 2. With this metric, an MI of 1 reflects responses that are maximally modulated - i.e., return completely to baseline after each inhalation, while an MI of 0 reflects no inhalation-linked modulation. For widefield OMP-GCaMP8m imaging data (Fig. 6), because of weak but widespread inhalation-linked modulation prior to odorant presentation, MI was defined as the standard deviation of the high-pass filtered ROI trace after the first second of odorant presentation, divided by that of the same trace during the pre-odor period. A high-pass filter cutoff of 1 Hz was used to eliminate tonic signal components. In this case, an MI index of 1 indicates no additional odorant-evoked modulation. MI values were computed on a trial-by-trial basis and averaged across repeated trials of odorant.

#### Mitral and tufted cell somatic signal analysis

To identify odorant-responsive mitral cells, imaging sequences were first movement-corrected in the XY plane using rigid registration, followed by somatic segmentation in Suite2P (Pachitariu et al., 2017). Initial segmentation was manually curated in the Suite2P environment, and the resulting ROIs were exported for further analysis in our custom Matlab Image Analysis GUIs. Daughter mitral cells innervating the parent glomerulus of interest were identified by their responsiveness to a near-threshold concentration of the diagnostic odorant, using an auROC (area under the receiver operating characteristic curve) criterion comparing the fluorescence signal during and prior to odorant presentation, compiled across 4 - 6 trials. In most cases, an auROC cutoff of 0.8 was used to identify responsive cells. For tufted cell analysis, automated segmentation was not used due to confounding signal from extensive lateral dendrites in the same focal plane. Instead, responsive cells were manually segmented based on ΔF response maps, and daughter cells identified by visually tracing their primary dendrites to the parent glomerulus. Following cell identification, for most analyses signals from all identified daughter mitral or tufted cells (range, 2 - 7 per glomerulus) were averaged for analysis of inhalation-linked dynamics, to improve signal-to-noise ratio.

#### Paired acid-ester comparisons

In most cases, comparison of response metrics between acids and their ester or aldehyde derivatives (e.g., Fig. 4) were made for odorant dilutions matched relative to their near-threshold concentration, and typically at 10x near-threshold unless specified otherwise in the Text. These values were consistent in different mice for the same odorants, and matched dilutions elicited responses of similar peak amplitude (± 30%). Net excitation ratio (e.g., Fig. 7O,P) was defined as the ratio of the mean non-negative activity in the first second of odorant presentation of the ester, divided by the same metric for the corresponding acid. GCaMP8f signals were made non-negative by subtracting the minimum ΔF/F value of either odorant response from both traces.

#### Statistical analysis

To compare response latencies between acids and their corresponding functional group derivatives (esters, aldehydes, alcohols), separate linear mixed-effects models were fit in R using the lme4 package (Bates et al., 2015) for each functional group comparison. Response latency was modeled as a function of functional group (acid versus derivative) and odorant family (tiglic, 2-methyl butyric, phenylacetic, isovaleric) as fixed effects, with biological replicate modeled as a random effect nested within odorant family to account for repeated measurements obtained from the same identified glomerulus across functional groups.

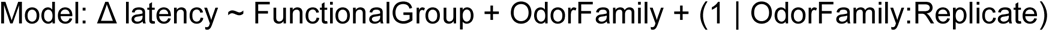

Estimated marginal means (emms) were calculated using the ‘emmeans’ package in R (https://rvlenth.github.io/emmeans/) and pairwise comparisons between acids and the corresponding functional group were performed for each model. For analyses involving multiple planned functional-group comparisons, P values were adjusted using the Bonferroni correction to control the family-wise error rate. For analyses involving a single planned comparison (e.g., acid versus ester), pairwise comparisons of estimated marginal means were performed without adjustment for multiple testing. To directly compare acid and ester response latencies, taking into account different odorants and biological replicates, we fit an analogous mixed-effects model and made pairwise comparisons of estimated marginal means without adjustment for multiple testing.

To compare relative response latencies in non-primary glomeruli for different odorants (Fig. 5E), latencies relative to the primary glomerulus, measured in the same trials, were analyzed using a one-way analysis of variance (ANOVA) implemented in R, with odorant identity as the independent variable. Odorant-specific emms (calculated using ‘emmeans’) were tested against a null value of zero to determine whether response latencies differed significantly from baseline, and Tukey-adjusted pairwise comparisons were used for post-hoc assessment of differences in latency between odorants.

To evaluate timing-based discrimination of ester and acid-evoked responses (Fig. 6 H, I, J), receiver operating characteristic (ROC) analysis was conducted in Python using the SciPy package. True labels were shuffled 5,000 times and the resulting ROC curves for MI and latency were pooled to create a combined null distribution by plotting the 2.5-97.5 percentiles.

## Acknowledgements

We thank Victoria Kent, Conor Craig, and Gustavo Vasquez for technical support; Crystal Davey and the University of Utah Transgenic Gene-Targeting Core for gene targeting design and mouse generation services; Alla Borisyuk and Parikshita Gya for assistance with CFD-PBPK model generation, and the CALIS network resources (doi: 10.15454/9PGN-W156).

## Funding

MW: NIH DC013076, NS109979, NSF NeuroNex 201427

MW/HM: DC022770

MAH: F32DC022517

JMH: ANR-18-CE92-0018-01

KZ: NIH-NIDCD R01 DC020737 and R01DC020302 to KZ

---

**Table S1.**
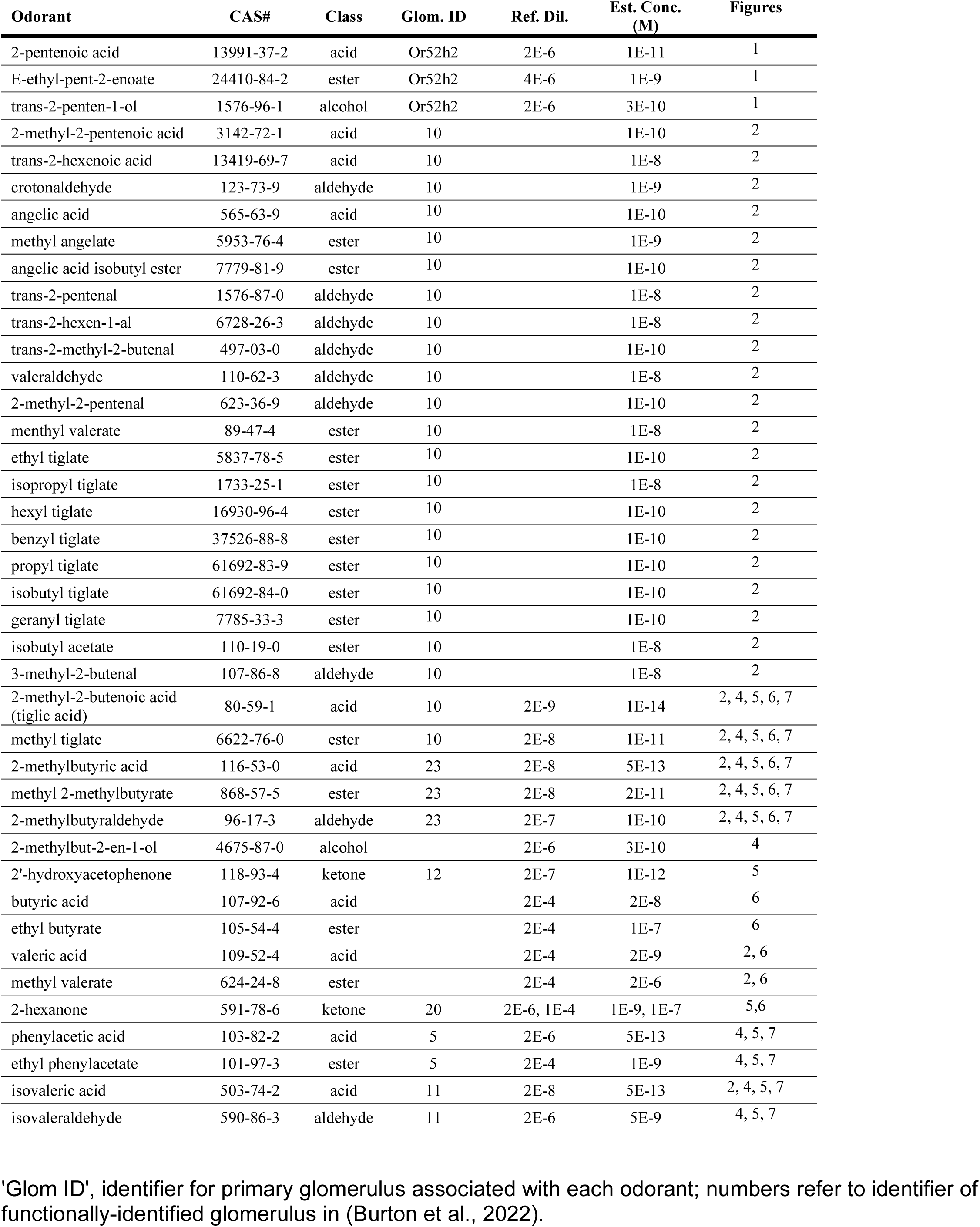
Odorants, reference dilutions and estimated concentrations used.

**Supplementary Figure S1 (related to Figure 1).**
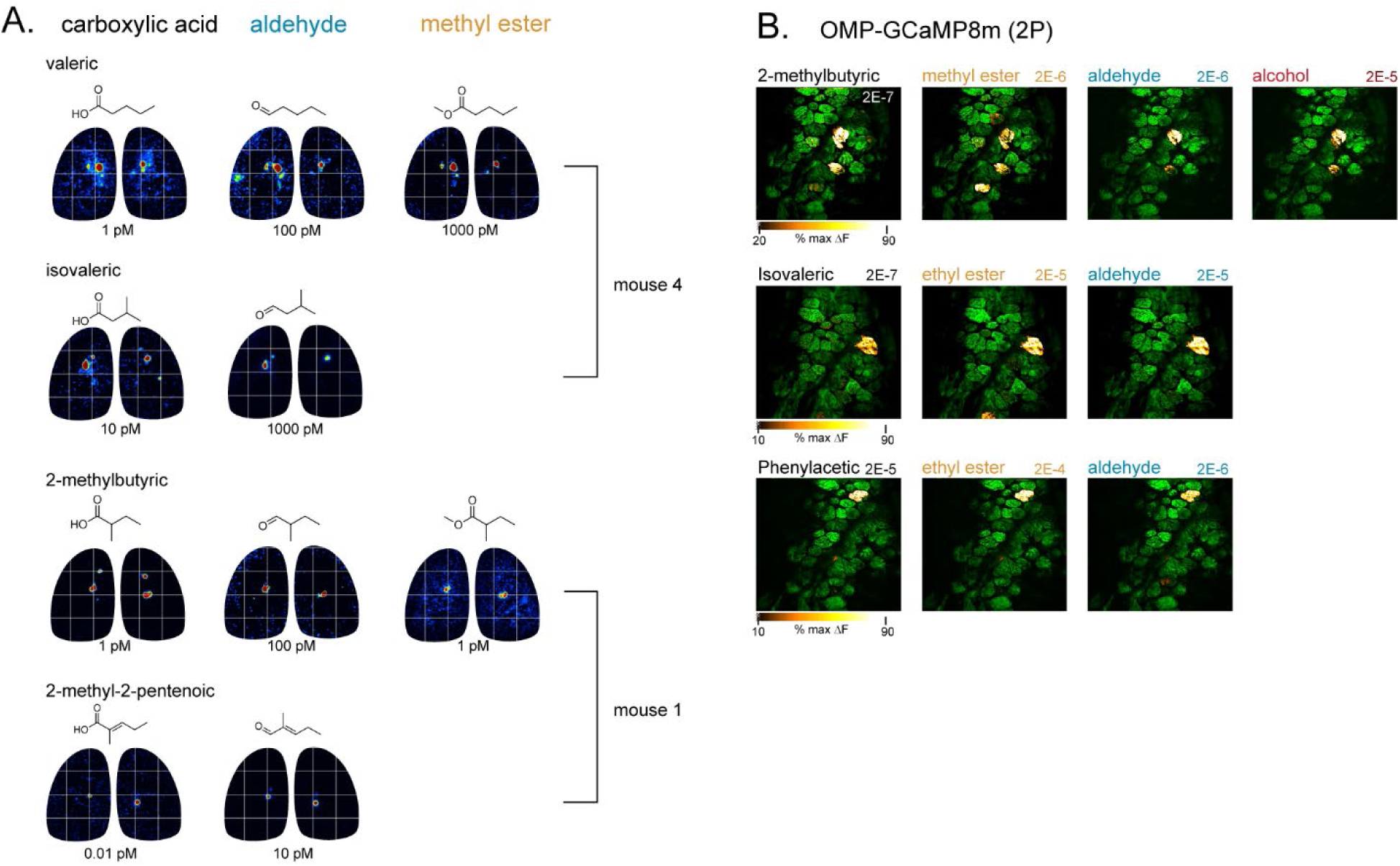
Potent co-tuning of functionally-identified glomeruli to ester, aldehyde and alcohol derivatives of the primary carboxylic acid. **A.** Epifluorescence response maps of OSN inputs (OMP-IRES-tTA x tetO-GCaMP6s) showing near-identical selective activation of dorsal glomeruli to four additional carboxylic acids and their ester and aldehyde derivatives, at estimated delivered concentrations shown. Data from (Burton et al., 2022). **B.** Two-photon response maps of OSN inputs (OMP-Cre x TIGRE2-GCaMP8m) to glomeruli identified by their sensitivity to 2-methyl butyric acid (2-MBA) and isovaleric acid (IVA), with co-tuning to ester, aldehyde and alcohol derivatives. Images taken from the same mouse (different from Fig. 1C).

**Supplementary Figure S2 (related to Figure 3).**
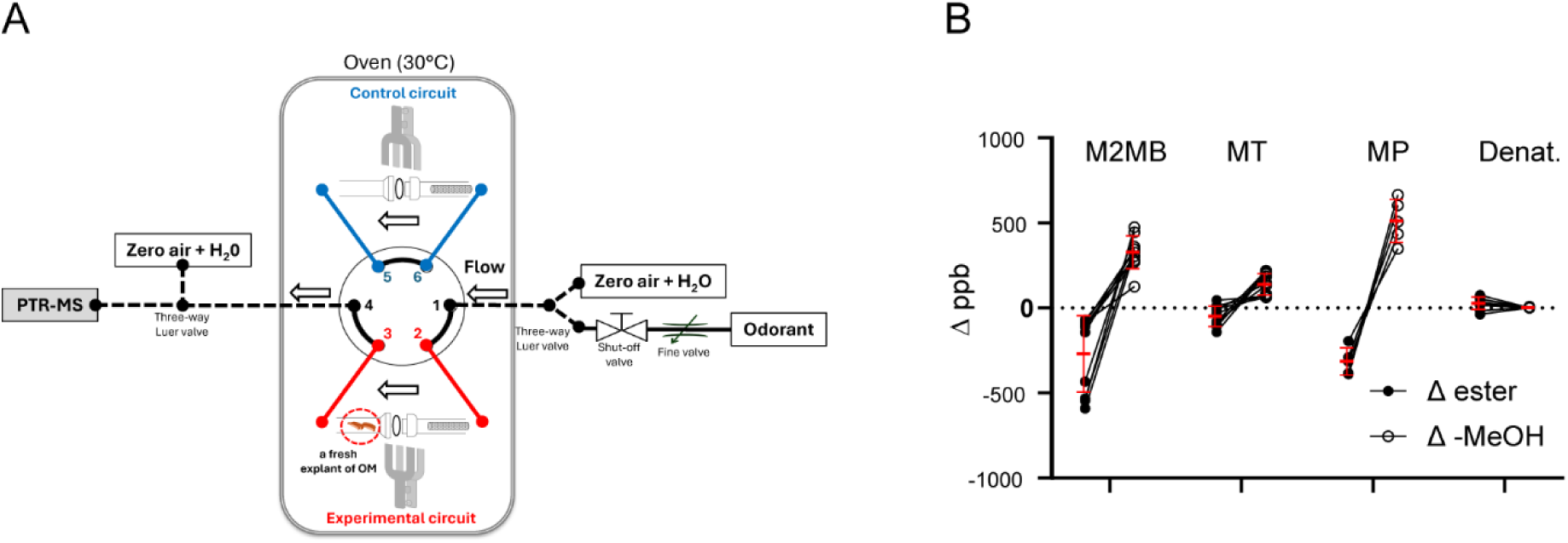
Details of PTR-MS measurement of odorant metabolism by mouse olfactory mucosa. **A.** Detailed schematic of setup for time-resolved measurement of ester metabolism by olfactory mucosa using PTR-MS. Air flow is from right to left and follows either a control circuit (blue) or an experimental circuit (red) containing the mucosa explant before travelling to the PTR-ToF-MS detector. **B.** Summary of reductions in ester signal amplitude and concomitant increases in methanol (MeOH) signal amplitude for olfactory mucosa flow, relative to control flow circuit, for different methyl esters tested. M2MB, methyl2-methyl butyrate; MT, methyl tiglate; MP, methyl propionate.

**Supplementary Fig. S3 (related to Figure 4).**
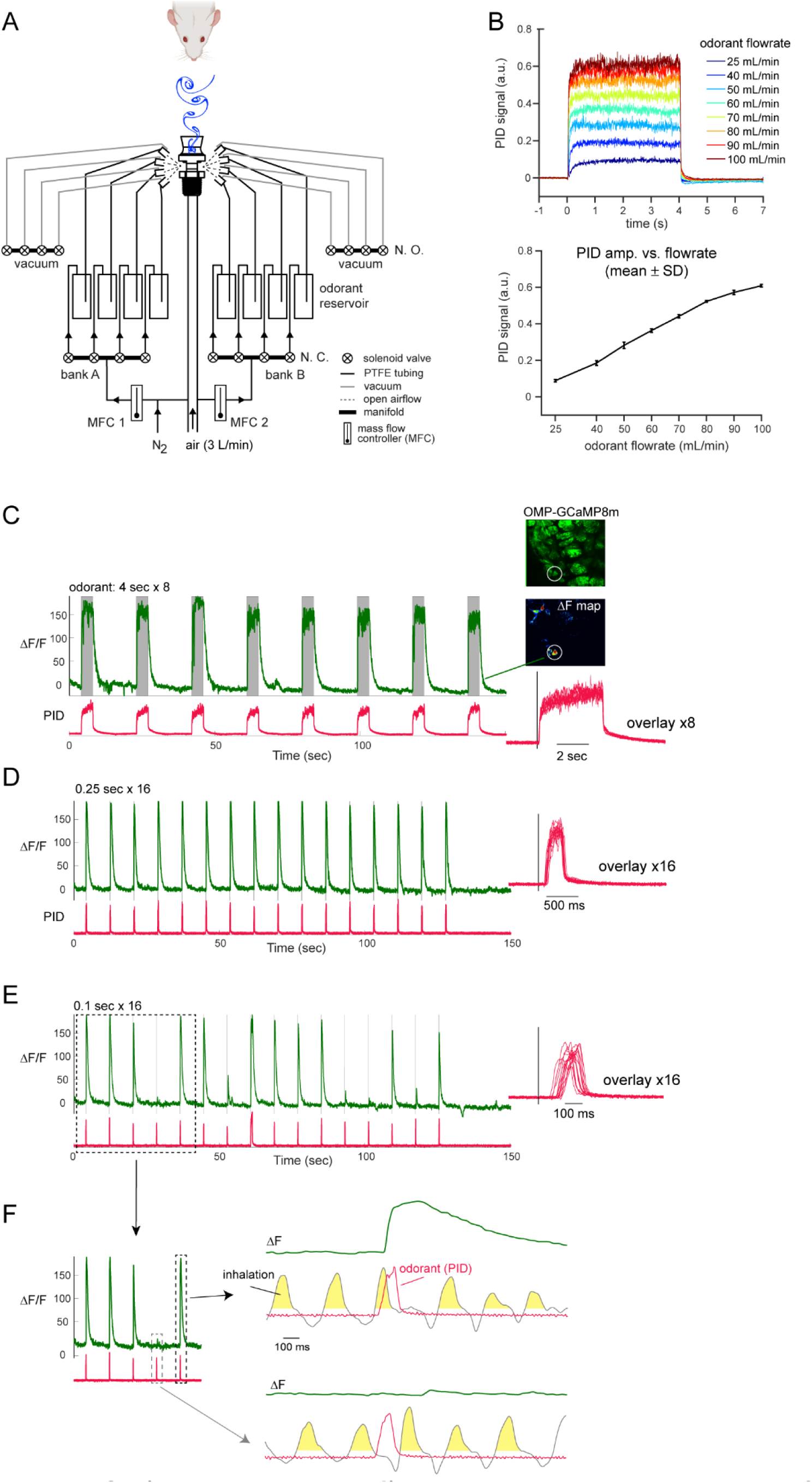
Design and characterization of olfactometer for control of odorant stimulus timing. **A.** Schematic of olfactometer design. **B.** Top: Photoionization detector (PID) traces of odorant concentration at different flowrates through odorant vial. Bottom: Mean ± SD. of PID signal amplitude versus odorant reservoir flowrate, showing a linear relationship over a four-fold range. **C.** PID traces (bottom) and OSN-GCaMP8m signals evoked by repeated 4-sec odorant presentations in the awake mouse. Top insets show mean F and ΔF images with imaged glomerulus. Lower inset shows overlay of PID traces from the 8 trials. **D.** Same as (C), showing consistent responses to 16 presentations of 0.25 sec duration pulse. **E.** Same as (D), with 16 presentations of 0.1 sec pulse. PID trace shows some loss of precision in timing, and OSN-GCaMP8m trace shows missed responses. **F.** Expansion of boxed region in (E), with respiration trace overlaid for individual presentations (right). Yellow-shaded areas indicate time period of inhalation. Note that ‘missed’ OSN response corresponds to odorant onset and offset occurring in between inhalations.

**Supplemental Figure S5. (related to Fig. 5).**
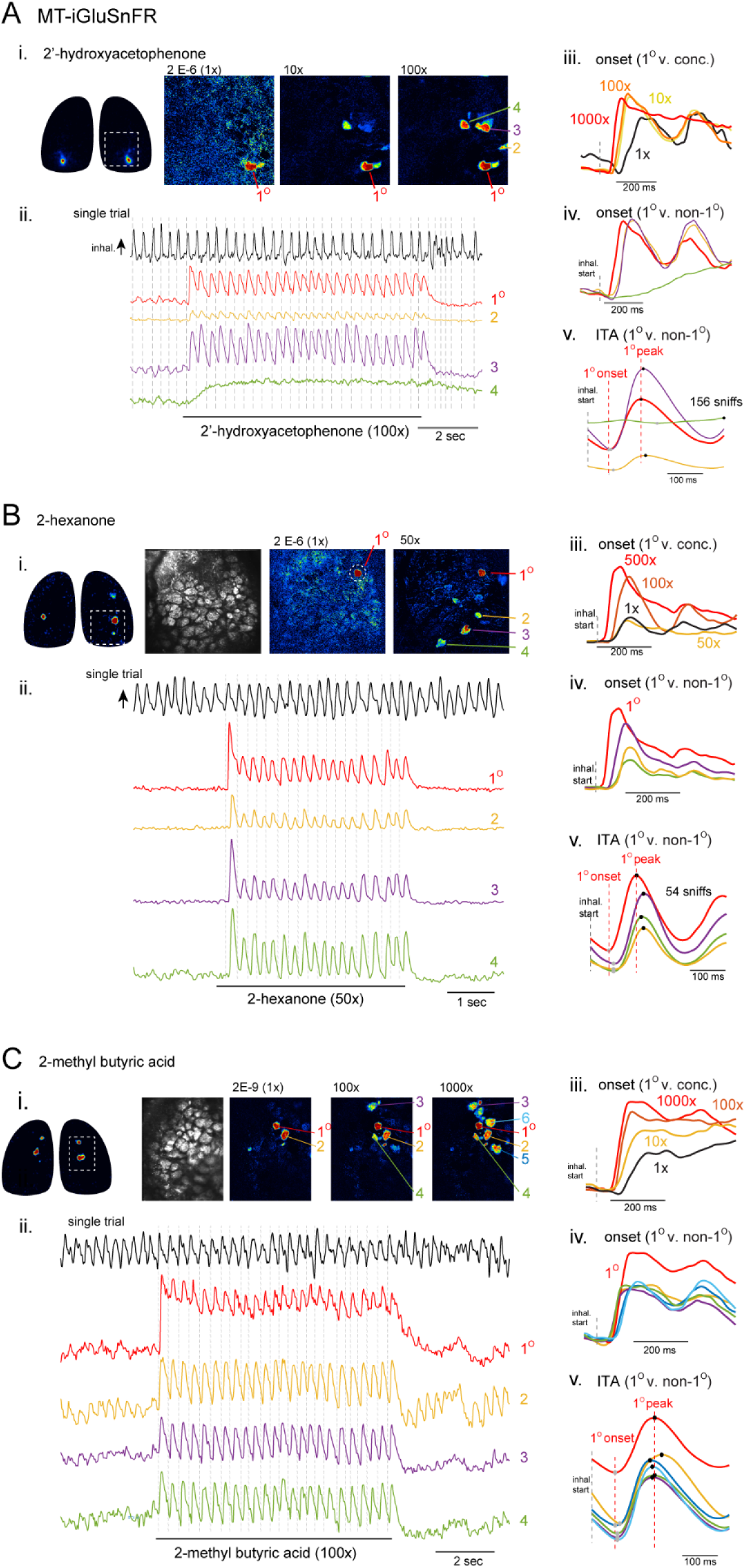
Sensitivity-timing relationships for additional functionally-identified glomeruli. A-C. Response maps and inhalation-linked timing for MT-iGluSnFR3 signals in primary (i.e., most-sensitive) and recruited (non-primary) glomeruli for 2-hexanone (A), 2’-OH acetophenone (B) and 2-methyl butyric acid (C), plotted as shown in Fig. 5B-D. (i) Widefield OMP-GCaMP6s response map showing singular activation, taken from Burton et al. (Burton et al., 2022). (ii) Single trial of odorant presentation with traces from primary and select non-primary glomeruli, with respiration trace (top) and inhalation times (dashed line). (iii) Inhalation-aligned response onset for primary glomerulus evoked by increasing odorant concentrations, referenced to near-threshold concentration (1x). (iv). Inhalation-aligned response onsets in primary and non-primary glomeruli evoked by suprathreshold odorant concentration. (v). Inhalation-triggered (ITA) traces generated from the same glomeruli and suprathreshold trials as in (iv).

**Supplementary Figure S6, related to Fig. 6.**
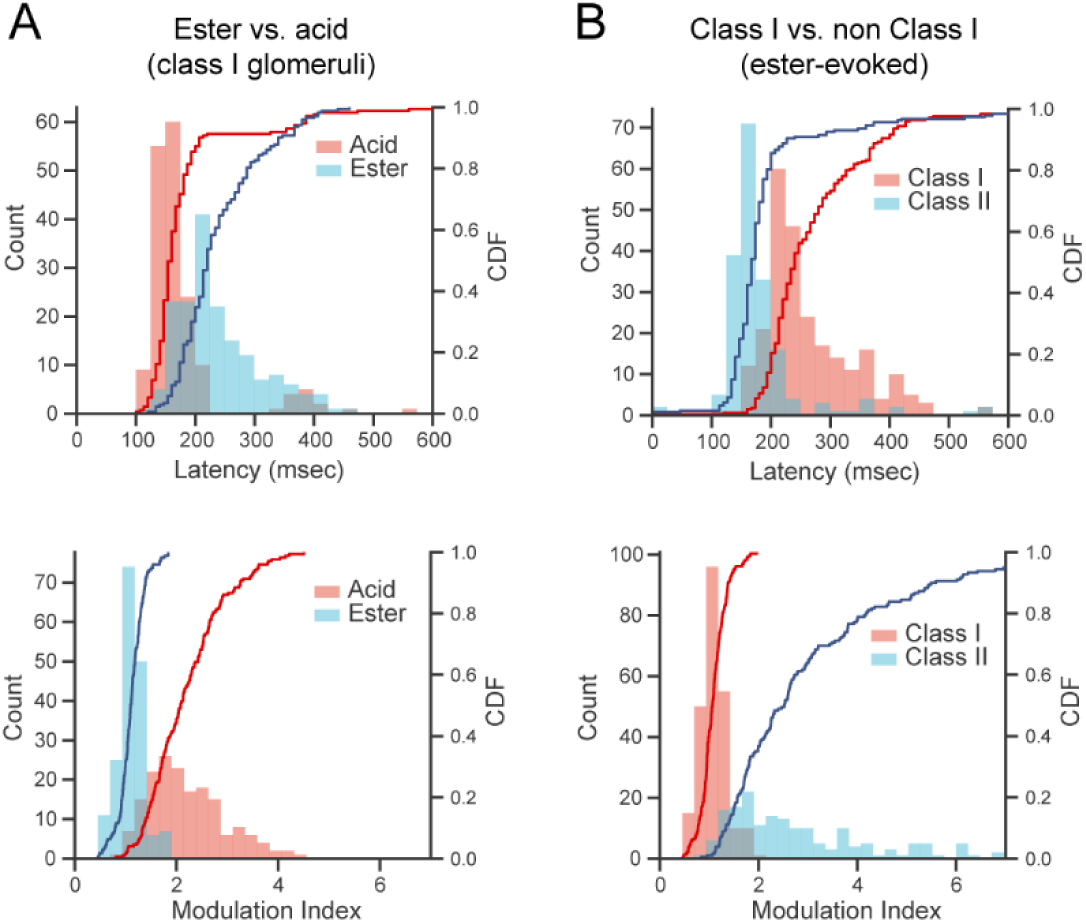
Distributions of onset latencies and modulation metrics distinguishing acid from ester-evoked responses and class I from class II glomeruli. **A.** Histograms and cumulative distribution function (CDF) plots for onset latency (top) and modulation index (bottom) for acid- and ester-evoked responses in co-tuned glomeruli; same data as in Fig. 6I. **B.** Histograms and CDF plots for onset latency and modulation index for ester-evoked responses in presumptive class I (acid-sensitive) and non-class I glomeruli; same data as in Fig. 6J.

**Supplementary Figure S7, Related to Figure 7.**
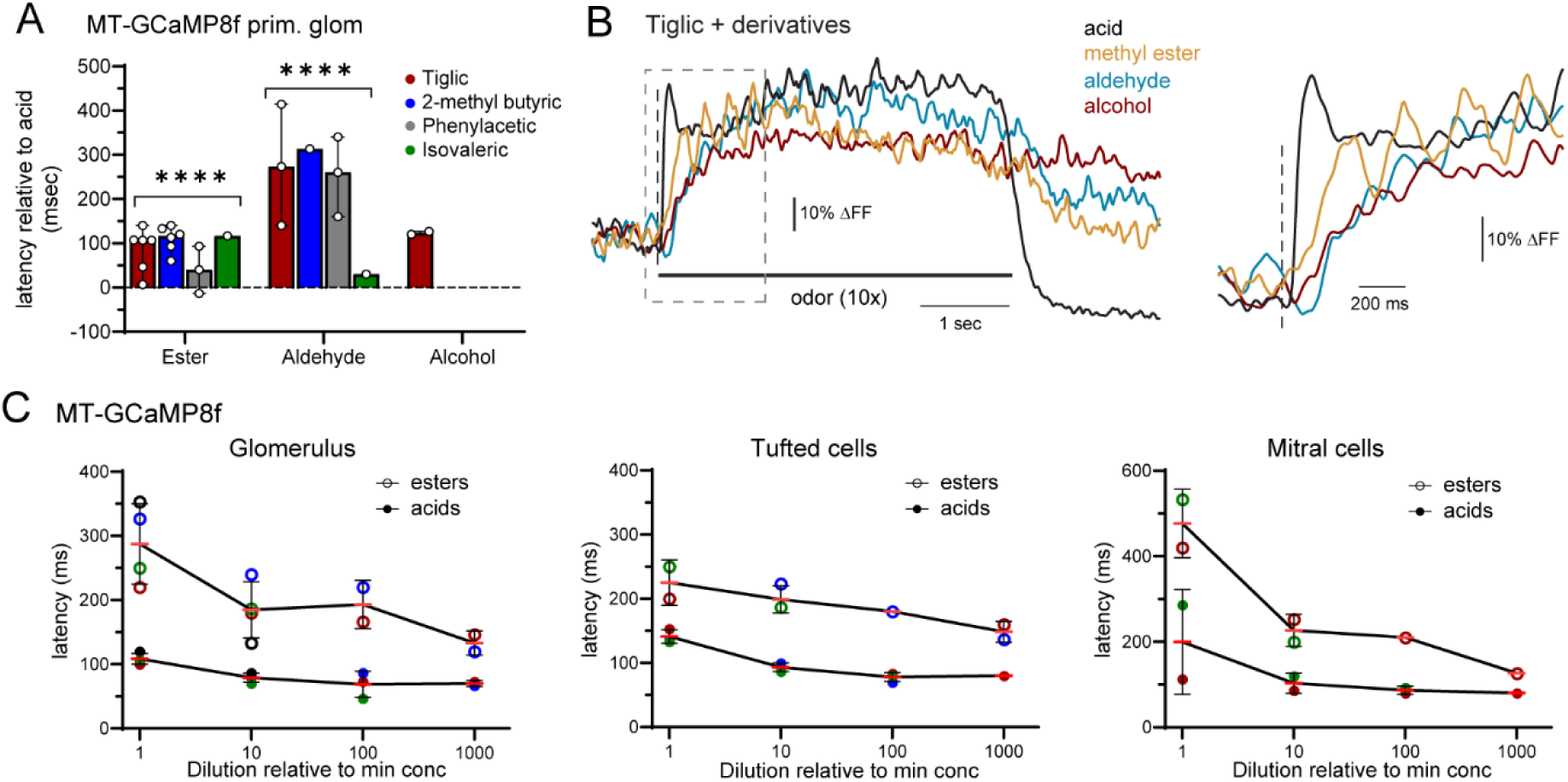
Delayed mitral and tufted cell responses to ester, aldehyde and alcohol derivatives relative to their primary carboxylic acid odorants. **A.** MT-GCaMP8f glomerular onset latency delays grouped by replicates of ester, aldehyde and alcohol derivatives, relative to their parent acid, plotted for each identified glomerulus. Points indicate biological replicates per glomerulus; bars indicate median, error bars indicate range. Same data as Fig. 7D. \*\*\*\**p* < 0.0001. Linear mixed-effects model followed by Bonferroni-corrected pairwise comparisons of estimated marginal means (EMMs) between acids and their corresponding esters and aldehydes. Alcohols not tested (insufficient n). **B.** Example traces from a tiglic acid-sensitive glomerulus showing inhalation-aligned MT-GCaMP8f response to acid, ester, aldehyde and alcohol derivatives. Right: Expansion of boxed area highlighting response onset. Odorants tested at ∼10x perithreshold dilution. **C.** MT-GCaMP8f onset latencies for acid-ester pairs as a function of concentration, referenced to near-threshold dilution, for signals in primary glomeruli (left), and their daughter tufted cells (middle) and mitral cells (right). Colored circles indicate median latency across biological replicates for a given acid or ester (same color-coding as in (A)). Red lines indicate mean of medians across acids or esters; error bars are standard deviation. All signals show longer latencies to the ester derivative at all dilutions.

**Supplementary Figure S8, Related to Figure 7.**
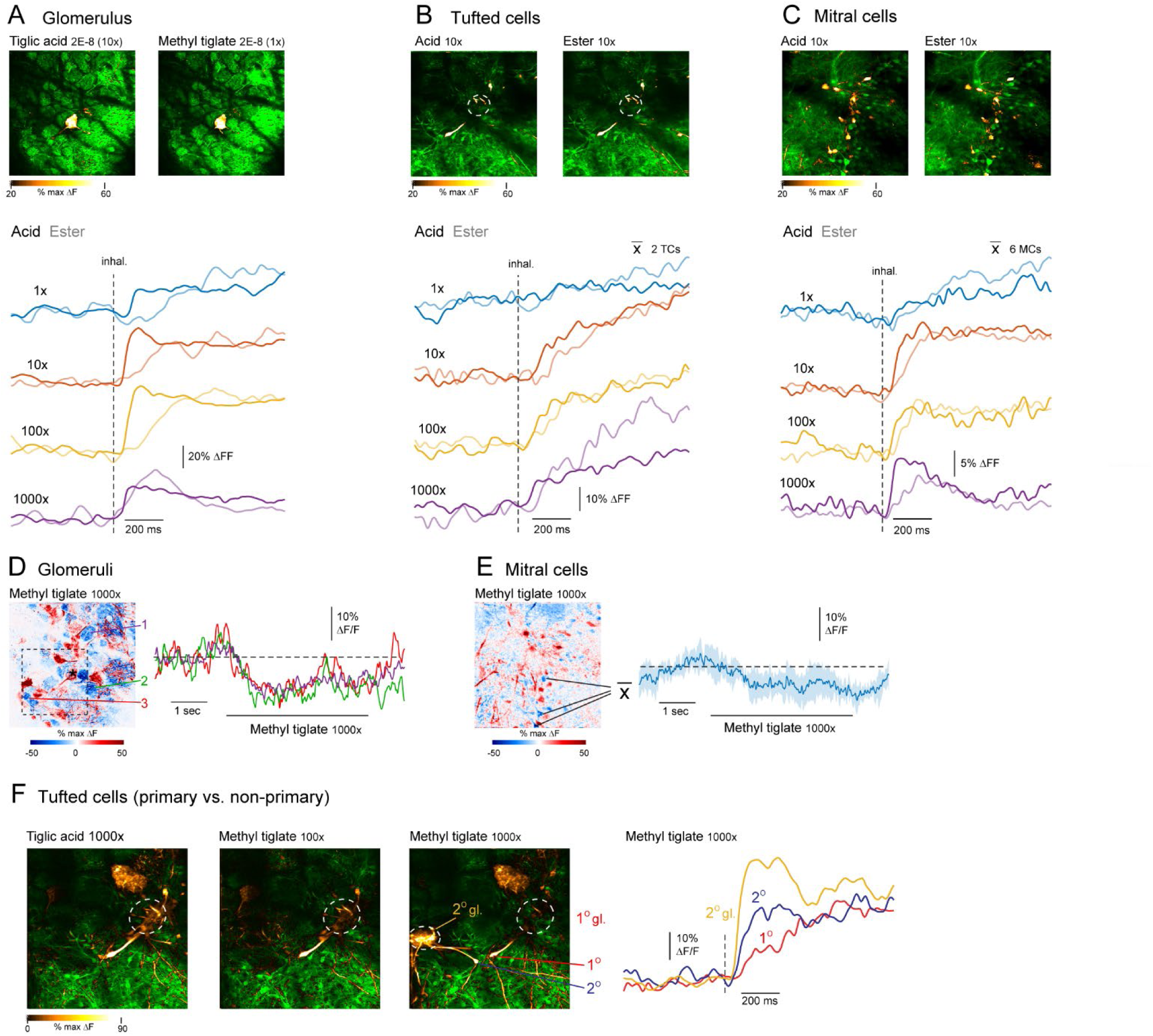
Delayed excitatory responses to ester derivatives persist across concentration despite suppression and faster excitation in neighboring glomeruli. A-C. Maps and inhalation-aligned traces of MT-GCaMP8f responses to tiglic acid or its methyl ester derivative methyl tiglate in the primary glomerulus (A) or its daughter tufted cells (B) or mitral cells (C). Top row: response maps to perithreshold concentrations of the acid and ester, showing singular activation of the primary glomerulus (A) and identifying excited daughter cells (B, C). Below are traces to the acid and ester, matched to the same concentration relative to near-threshold (1x) are scaled as per the scale bar. All traces in a sub-panel are shown on the same absolute scale (ΔF/F). Glomerular and mitral cell signals show reduced-amplitude response at 1000x concentration but slower excitatory response to the ester persists. Traaces in (B) and (C) are averages across the same 2 and 6 daughter cells, respectively. **D.** Left: GCaMP8f response map from the glomerular layer (same field of view as (A)) scaled and pseudocolored to show excitatory (red) and suppressive responses (blue) to 1000x methyl tiglate. Asterisk indicates primary glomerulus shown in (A). Note additional excited (non-primary) glomeruli as well as neighboring suppressed glomeruli. Right: Traces from three glomeruli showing suppressive responses. **E.** Response map and traces from the mitral cell layer (same field of view as (C)) revealing suppressive responses in neighboring mitral cells. Trace at right shows mean signal from three cells. **F.** Response maps and traces showing glomeruli and daughter tufted cells, imaged from region outlined in (D) in response to suprathreshold concentrations of tiglic acid or methyl tiglate. Primary tiglic acid/methyl tiglate glomerulus is outlined in white; note clear innervation by daughter tufted cell. Maps sequence shows recruitment of new glomerulus with concentration increase from 100x to 1000x methyl tiglate, with daughter tufted cell clearly apparent (right map). Right: Traces taken from the adjacent daughter tufted cells innervating the primary and non-primary glomeruli (labelled 1’ and 2’, respectively), and the parent non-primary glomerulus (’2’ gl’), in response to 1000x methyl tiglate. Note that secondary glomerulus responds with shorter latency than primary, and latency differences persist in their respective daughter tufted cells.

## Notes

### Competing Interest Statement

The authors have declared no competing interest.

